# An engineered transcriptional reporter of protein localization identifies regulators of mitochondrial and ER membrane protein trafficking in high-throughput screens

**DOI:** 10.1101/2021.04.11.439362

**Authors:** Robert Coukos, David Yao, Mateo I. Sanchez, Eric T. Strand, Jonathan S. Weissman, Michael C. Bassik, Alice Y. Ting

## Abstract

The trafficking of specific protein cohorts to the correct subcellular location at the correct time is essential for every signaling and regulatory process in biology. Gene perturbation screens could provide a powerful approach to probe the molecular mechanisms of protein trafficking, but only if protein localization or mislocalization can be tied to a simple and robust phenotype for cell selection, such as cell proliferation or FACS. To broadly empower the study of protein trafficking processes with gene perturbation, we developed a genetically-encoded molecular tool named HiLITR. HiLITR converts protein colocalization into proteolytic release of a membrane-anchored transcription factor, which drives the expression of a chosen reporter gene. Using HiLITR in combination with FACS-based CRISPRi screening in human cell lines, we identify genes that influence the trafficking of mitochondrial and ER tail-anchored proteins. We show that loss of the SUMO E1 component SAE1 results in the mislocalization and destabilization of mitochondrial tail-anchored proteins. We also demonstrate a distinct regulatory role for EMC10 in the ER membrane complex, opposing the transmembrane-domain insertion activity of the complex. Through transcriptional integration of complex cellular functions, HiLITR expands the scope of biological processes that can be studied by genetic perturbation screening technologies.

## Introduction

Gene perturbation screens, in which libraries of cells bearing individual genetic perturbations are assessed for fitness, growth, or other phenotypes, have been broadly applied to uncover the genetic bases of specific cellular processes. For example, CRISPR-based knockout screens have identified essential human genes (Hart et al., 2015; Shalem et al., 2014; Wang et al., 2015), discovered factors that confer drug or toxin resistance (Wang et al., 2014; Zhou et al., 2014), and dissected signaling and regulatory networks (Klann et al., 2017; Parnas et al., 2015). In general, gene perturbation screens can be implemented in either pooled or arrayed formats. Pooled screens simplify the handling of large libraries with >10^5^ unique elements (Kampmann et al., 2015; Morgens et al., 2016; Wang et al., 2015), but require that the cellular function of interest be coupled to a simple readout, such as cell proliferation (Han et al., 2020; Kory et al., 2018) or FACS (Dejesus et al., 2016; Potting et al., 2017). Arrayed screens, on the other hand, are well-suited to complex readouts, such as high-content or time-lapse microscopy, and have been used to discover factors that regulate cell division (Neumann et al., 2010), endocytosis (Liberali et al., 2014), and membrane protein trafficking (Hansen et al., 2018; Krumpe et al., 2012). However, compared to pooled screens, arrayed screens are usually noisier, limited to smaller libraries (10^3^-10^4^), and more technically difficult and time-consuming to implement, requiring specialized instrumentation not available to all laboratories.

To combine the strengths of the pooled screen format (library size, simplicity) and the arrayed screen format (versatility in readout), we sought to develop a molecular reporter capable of converting complex cellular processes such as protein trafficking or mislocalization into simple single-timepoint, intensity-based FACS readouts. Such a tool would enable screening of large libraries in a pooled format without sacrificing the versatility and specificity required to probe more complex cellular processes.

Here, we report HiLITR (High-throughput Localization Indicator with Transcriptional Readout), a molecular tool that converts protein localization or mislocalization into simple expression of a fluorescent protein. We used HiLITR in combination with a human CRISPRi library to screen for factors that regulate the trafficking of mitochondrial and ER membrane proteins. We found that knockdown of the SUMO E1 ligase component SAE1 selectively destabilizes and increases the mislocalization of mitochondrial tail-anchored proteins. Additionally, we found that knockdown of the EMC10 subunit of the ER membrane complex (EMC) increases insertion of specific tail-anchored proteins into the ER membrane, in opposition to the function of other EMC components.

### HiLITR is a live-cell transcriptional reporter of protein localization

To design HiLITR, we required a mechanism to convert protein localization or mislocalization in live cells to a simple readout for pooled genetic screens. To do so, we designed two protein components – a protease (GFP-TEVp) and a transcription factor (TF), each targetable to specific subcellular locations (Fig. 1A/B). If the protease and TF are colocalized to the same organelle (e.g. the mitochondrial membrane in Fig. 1A), then proximity-dependent proteolysis of a protease cleavage sequence (TEVcs) in the TF’s anchor releases the TF, which subsequently translocates to the nucleus and drives expression of a chosen reporter gene. If the protease and TF are not co-localized, then TF cleavage and reporter gene expression will not occur.

**Figure 1.**
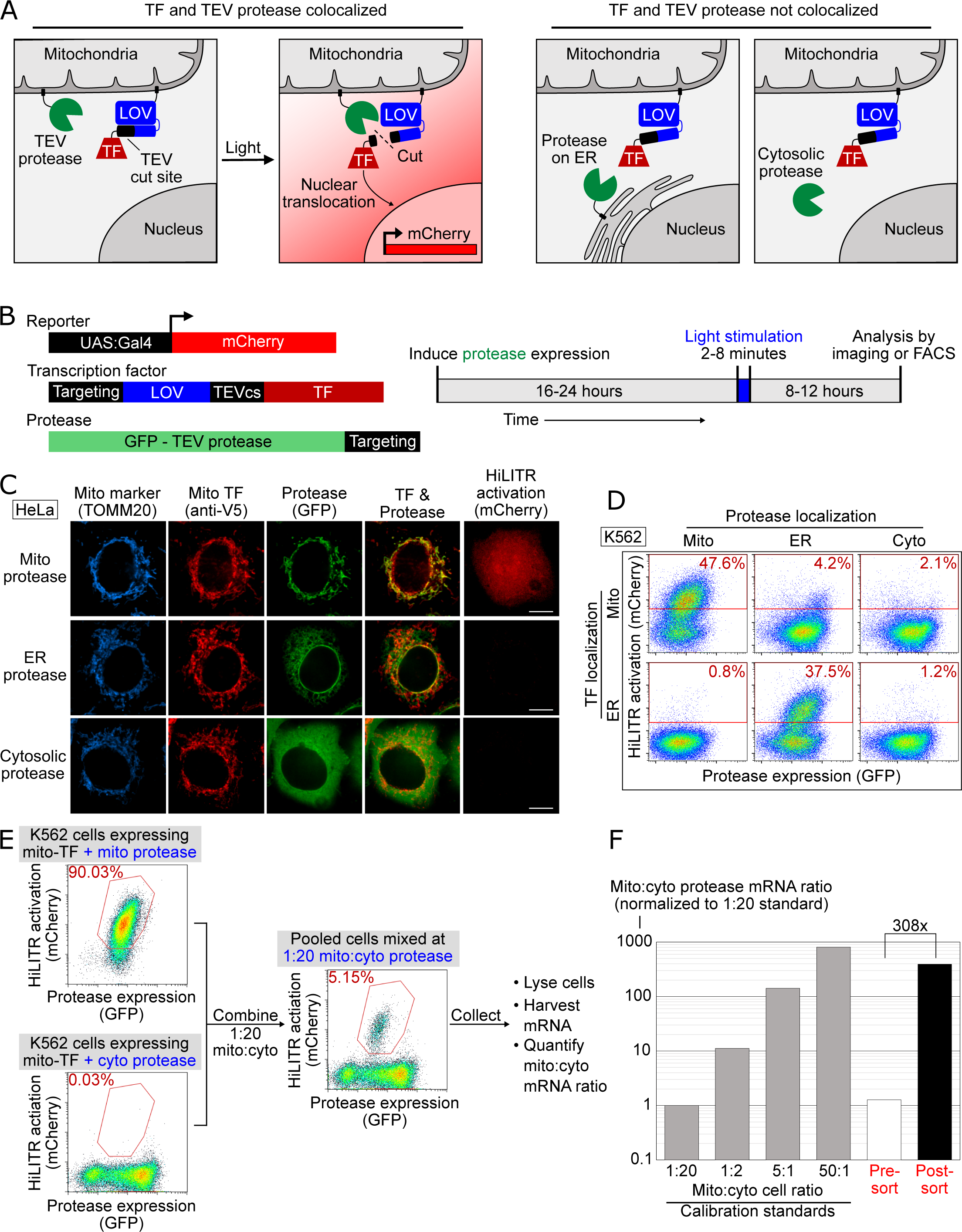
HiLITR provides transcriptional readout of protein localization in live cells. **(A)** Schematic of HiLITR. HiLITR has two components: a low-affinity protease (green) and a membrane-anchored transcription factor (TF, red). Left: when protease and TF are colocalized on the same organelle, and 450 nm blue light is supplied, the TF is released by proximity-dependent cleavage and drives reporter gene expression. Right: when protease and TF are not colocalized, HiLITR is off. **(B)** Domain structures of HiLITR components and timeline of HiLITR. The targeting domain is a protein or localization peptide that directs the TF /protease to the desired subcellular region (such as the mitochondrion in (A), left). **(C)** Fluorescence images of HiLITR in HeLa cells. TF is on the outer mitochondrial membrane (OMM), and protease is localized to the OMM (top row), ER membrane (middle), or cytosol (bottom). mCherry is the reporter gene and TOMM20 is a mitochondrial marker. Cells were stimulated with 450 nm light for 3 minutes. Scale bar, 10 µm. **(D)** FACS plots of K562 cells expressing HiLITR. TF is on the OMM (top row) or ER membrane (bottom row), while protease localization is varied as indicated. Light stimulation was 3 min. mCherry on the y-axis reports HiLITR activation, and GFP on the x-axis reports protease expression level. Percentage of cells above the red line is quantified in each plot. **(E)** Model selection on K562 cells expressing HiLITR TF on mitochondria. Cells with mitochondrial protease (co-localized with TF) versus cytosolic protease (not co-localized with TF) were combined in a 1:20 ratio. Cells were stimulated with light for 3.5 minutes and sorted for high mCherry expression 8 hours later. **(F)** qPCR analysis of mito- and cyto-protease transcript from predefined, pre-sort, and post-sort cell mixtures from (E). Mito-protease cells were enriched 308-fold over cyto-protease cells in one round of sorting.

To maximize the dynamic range of HiLITR, we included a second “gate” in our design, in the form of a photosensory LOV domain adjacent to the TEVcs in the TF tether (Figure 1B). The LOV domain sterically impedes protease cleavage in the dark, but provides access under blue light illumination. HiLITR therefore acts as an AND gate, requiring both protease/TF colocalization *and* blue light to turn on. This two-gate design improves dynamic range, temporal precision, and tunability compared to a single-input design (Kim et al., 2017).

To test HiLITR, we generated a TF construct targeting the outer mitochondrial membrane (OMM), and protease constructs targeting the OMM, ER membrane (ERM), or cytosol (Fig. S1). Transient transfection produced high background signal (Fig. S2A/B), which was alleviated by stable integration (Figs. S2C/D, S4A). We improved specificity by replacing TEV protease with the ∼5-fold more catalytically efficient “ultraTEV” (S153N) (Sanchez and Ting, 2020) and the eLOV domain with the tighter-dark state hLOV (Kim et al., 2017) (Fig. S2C-F). Replacement of the Gal4 TF with a more active Gal4-VP64 fusion also improved signal (Fig. S2G/H). Using the best HiLITR constructs, we then varied tool expression time, stimulation conditions, and reporter expression time to optimize the conditions for HiLITR use in multiple cell types (Fig. S3; Text S3; Fig. 1C)

We further tested the versatility of HiLITR by designing a TF targeted to the ER membrane. By FACS in K562 cells (Fig. 1D), we observed clear activation of ERM-TF by ERM-protease, but not by OMM-protease or cytosolic-protease. We also saw clear activation of OMM-TF by OMM-protease but not by ERM-protease or cytosolic-protease. The absence of cross-reactivity is striking given that mitochondria and ER form contacts in mammalian cells (Wu et al., 2018). Perhaps the fraction of HiLITR TF localized to these contact sites is very small compared to the total amount on the OMM or ERM surface, such that the contribution of ER-mito contacts to HiLITR activation is insignificant. As an additional test of specificity, we designed HiLITR constructs localized to the peroxisomal membrane, which also gave expected patterns of activation (Fig. S4F/G; Text S4A).

Finally, we performed a model selection to enrich cells with colocalized HiLITR components from cells with non-colocalized HiLITR components. qPCR analysis showed >300 fold enrichment in a single round of FACS (Figs. 1E/F, S5).

### Using HiLITR in pooled CRISPRi screens to probe pathways of ER and mitochondrial membrane protein trafficking

Because HiLITR provides a simple, fluorescence intensity-based readout of protein colocalization in living cells, we sought to combine it with CRISPRi and FACS in a pooled screen to identify factors regulating the trafficking of ER and mitochondrial membrane proteins. For instance, if the HiLITR TF is targeted to the OMM via an N-terminal transmembrane anchor (signal anchor) and the HiLITR protease is localized to the OMM via a C-terminal transmembrane anchor (tail anchor), then any sgRNA that disrupts a gene important for tail anchored OMM protein targeting should reduce HiLITR-driven reporter gene expression (Fig. 2A). Cells with reduced HiLITR activation can be enriched by FACS and analyzed by sgRNA sequencing (Fig. 2B).

**Figure 2.**
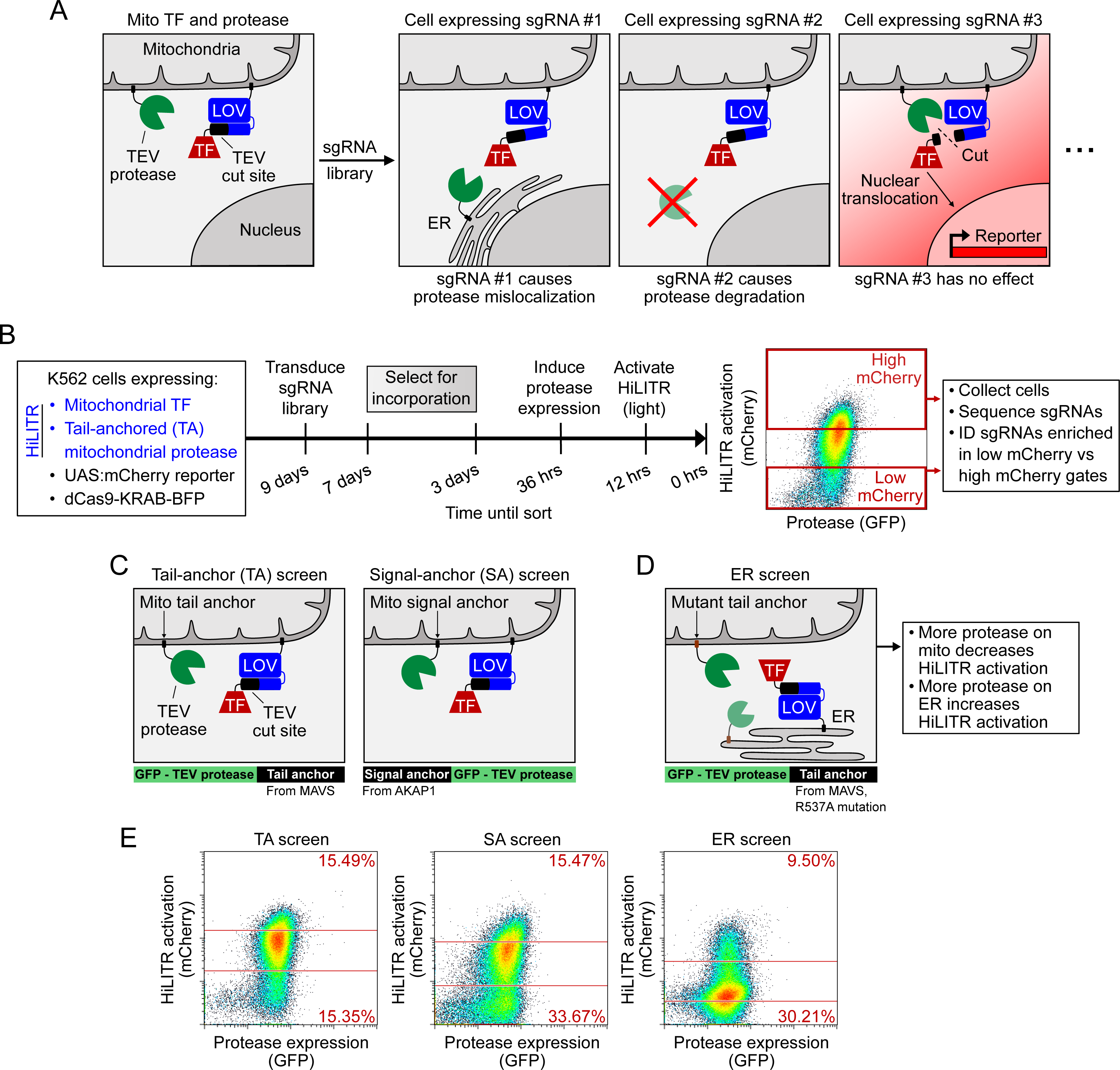
HiLITR reads out protein mislocalization or loss in CRISPRi screens. **(A)** Possible outcomes for sgRNA disruption of mitochondrial protease in cells expressing mitochondrial HiLITR TF and protease. Example sgRNA #1 disrupts protease localization while sgRNA #2 reduces protease abundance. Both perturbations lead to decreased HiLITR-driven mCherry expression. **(B)** Format and timeline of CRISPRi screens. **(C-D)** Three HiLITR cell lines used for CRISPRi screens. The first two (C) use mitochondria-localized TF and either tail-anchored (left) or signal-anchored (right) mitochondrial protease. The third cell line (D) uses ER-localized TF and a mutated tail-anchored mitochondrial protease. **(E)** FACS plots showing cell populations collected and sequenced from the TA, SA, and ER CRISPRi screens from C-D. Light stimulation times varied from 3.5-5 minutes.

A myriad of proteins function at the OMM and ERM, localized via signal anchored, tail anchored, internal, and multipass transmembrane domains. Several distinct pathways orchestrate the co-translational or post-translational insertion of these proteins at the ERM (Shao and Hegde, 2011) and at the OMM (Hansen and Herrmann, 2019). Recent studies have also revealed a striking interplay between ER and mitochondrial membrane targeting pathways (Costa et al., 2018; Gamerdinger et al., 2015; Mårtensson et al., 2019), such as the ER-SURF pathway, in which some OMM proteins are harbored on the ERM when mitochondrial import is taxed (Hansen et al., 2018).

Despite our detailed and evolving picture of ER and mitochondrial protein trafficking pathways, some major gaps in understanding remain. For example, the mechanisms by which mitochondrial proteins with C-terminal transmembrane domains (“tail anchors”) are delivered and inserted into the OMM are unclear. There are about 40 human mitochondrial tail-anchored (TA) proteins, including the apoptosis regulators BAX and BCL2 (Wolter et al., 1997) and mitochondrial fission proteins FIS1 (Yoon et al., 2003) and MFF (Gandre-Babbe and Van Der Bliek, 2008). The targeting of TA proteins presents a unique challenge because their hydrophobic transmembrane domains are translated last, and must be handed off from the ribosome to appropriate chaperones or lipids before aggregation or misfolding can occur. It is known that the Get/TRC pathway, responsible for targeting of ER tail-anchored proteins, is not required for targeting of mito TA proteins (Borgese et al., 2019). In yeast, the HSP70 protein STI1 (human STIP1) and Pex19 may play a role in mito TA protein targeting (Cichocki et al., 2018), but in vitro experiments with proteolytically shaved mitochondria show that unassisted or minimally-assisted insertion may also be possible (Kemper et al., 2008; Setoguchi et al., 2006).

We envisioned using three distinct HiLITR configurations to investigate mitochondrial tail-anchored protein insertion. In the most direct approach, impaired mitochondrial localization of a TA protease would reduce the release of a signal-anchored (SA) mitochondrial TF, diminishing HiLITR activation (“TA screen”, Fig. 2C and S1).

However, this “loss of signal” design may enrich false positive genes whose knockdown nonspecifically disrupts HiLITR component expression (e.g. ribosome proteins). To filter out such genes, we designed a second HiLITR screen with the same TF but a signal-anchored—rather than tail-anchored—protease (“SA screen”, Figs. 2C, S1). Comparison of TA and SA screen results should eliminate most false positives and identify factors that selectively influence the targeting of mito TA proteins.

For our third HiLITR configuration, we designed a “gain of signal” screen based on the hypothesis that disrupted or non-targeted mitochondrial TA protease may reroute to ER or Golgi membranes. Thus, we expressed an ER/Golgi-targeted HiLITR TF together with a mutated mitochondrial TA protease (mTA*) that has increased propensity for mislocalization to ER/Golgi membranes (“ER screen”, Figs. 2D, S1, S4D/E, and Text S4B). Given the partial ER localization of this mTA* protease, we anticipated that our ER screen might also help to identify factors that regulate the targeting of native ERM proteins.

To first narrow our list of candidate genes, we performed a whole-genome “TA screen” with the HiLITR components shown in Fig. 2C. Human K562 cells expressing HiLITR and dCas9-BFP-KRAB were transduced with a library of sgRNAs targeting 20,000 genes, at 5 guides/gene, with 1900 nontargeting controls. Using FACS, we collected cell populations with high mCherry expression (i.e. strong HiLITR activation) and low mCherry expression (Fig. S6A). Next-generation sequencing of sgRNA abundance in the collected populations indicated enrichment of genes related to protein folding, membrane trafficking, and proteasome function in the low-mCherry cell population (Fig. S6A/B, Text S6). As expected, we also enriched a number of gene expression-related false-positives, including ribosome subunits, mediator complex subunits, and mRNA binding proteins.

Using these results, we designed an sgRNA sublibrary (Table S2) for simultaneous screening in the three HiLITR configurations (TA screen, SA screen, and ER screen). In total, we transduced our three clonal HiLITR K562 cell lines with 2,930 sgRNAs targeting 586 genes (5 guides/gene) plus 500 nontargeting controls. The screens were performed as shown in Figure 2B, with infection, passaging, and sorting carried out at 2,000-10,000x coverage—a higher level than standard—in order to detect subtle or partial effects. For each screen, we collected cell populations corresponding to high mCherry reporter expression and low mCherry reporter expression (Fig. 2E).

### CRISPRi screens with three HiLITR configurations identify proteins that influence the localization of mitochondrial and ER membrane proteins

The combined results from our three HiLITR screens are shown in Figure 3A. Sequencing data were analyzed by CasTLE (Morgens et al., 2016), which assigns to each gene an effect size and an associated CasTLE score (a measure of significance, signed for effect direction). CasTLE scores in the TA and SA screens largely were concordant, reflecting the similarity between the two configurations (R^2^ = 0.69, compared to R^2^ = 0.28 for TA vs ER screens and 0.37 for SA vs ER screens). Out of 586 genes, sgRNAs against 270 of them impacted HiLITR turn-on significantly (at 10% FDR) in at least one screen (186 genes in 2+ screens). The TA screen and whole-genome screens used the same HiLITR configuration, and of the 50 most significant genes from the whole-genome screen that were included in the three-screen sublibrary, 47 were significant in the TA screen (at 10% FDR).

**Figure 3.**
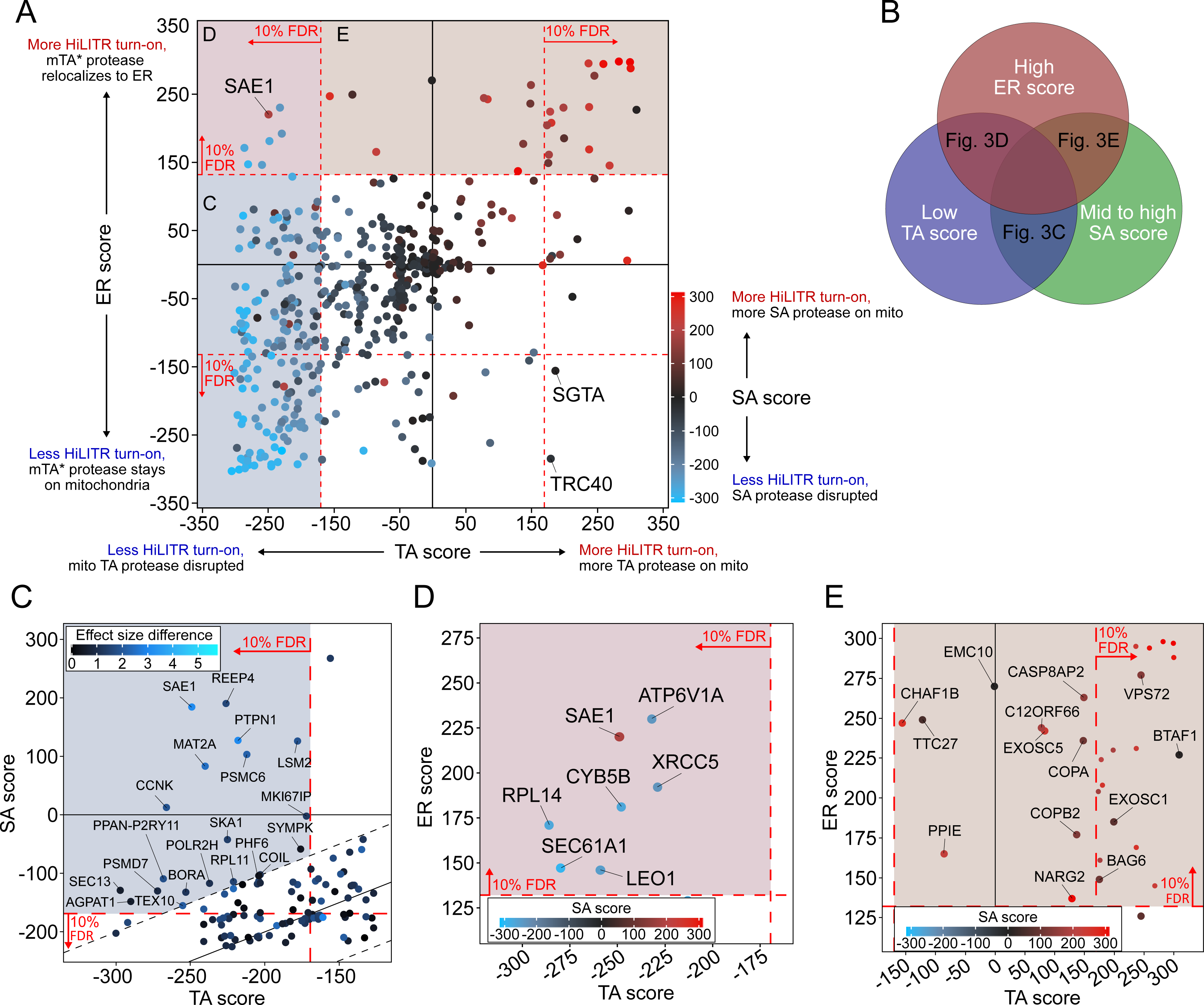
CRISPRi screen with HiLITR readout identifies proteins that influence the localization of mitochondrial membrane and ER membrane proteins. **(A)** CasTLE plot showing combined results from TA, SA, and ER CRISPRi screens. X-axis plots the TA screen score (lower when the mito TA protease from Figure 2C is disrupted) and y-axis plots the ER screen score (higher when the mito mTA * protease from Figure 2D relocalizes to the ER membrane). Points are color-coded according to score in the SA screen, with red denoting less disruption of the mito SA protease from Figure 2C. **(B)** Venn diagram showing that proteins regulating the targeting of mitochondrial tail-anchored proteins may exhibit some combination of low TA score, mid to high SA score, and high ER score. **(C)** Zoom-in of proteins with low TA score and medium-high SA score (replotted from (A)). Points are colored according to absolute difference in effect size in TA vs. SA screen. Dashed black lines enclose the 90% interquantile range for difference between TA and SA score. **(D)** Zoom-in of proteins with low TA score and high ER score, corresponding to maroon shaded region in (A). **(E)** Zoom-in of proteins with high ER score and medium-high SA score, corresponding to brown shaded region in (A). Unlabeled points showed significant increases in HiLITR activation (at 10% FDR) in all three screens and are likely to be nonspecific hits.

To assess the validity of our screens, we first checked genes with known roles in mitochondrial and ER protein trafficking. The TRC pathway (Schuldiner et al., 2008; Stefanovic and Hegde, 2007), which handles the membrane insertion of ER TA proteins, also mishandles overexpressed mitochondrial TA proteins (Vitali et al., 2018). Consistent with this activity, knockdown of the TRC pathway chaperones SGTA and TRC40 significantly altered HiLITR activation in the TA and ER screens, but not the SA screen (Fig. 3A; Text S7). Other TRC pathway components also altered HiLITR activation in the TA and ER screens (Fig S7; Text S7). The ER membrane complex (EMC), similar to the TRC pathway, handles insertion of a subset of TA proteins at the ERM (Guna et al., 2018). Among 9 subunits tested, 8 significantly altered HiLITR activation uniquely in the ER screen (Fig. 5A).

**Figure 4.**
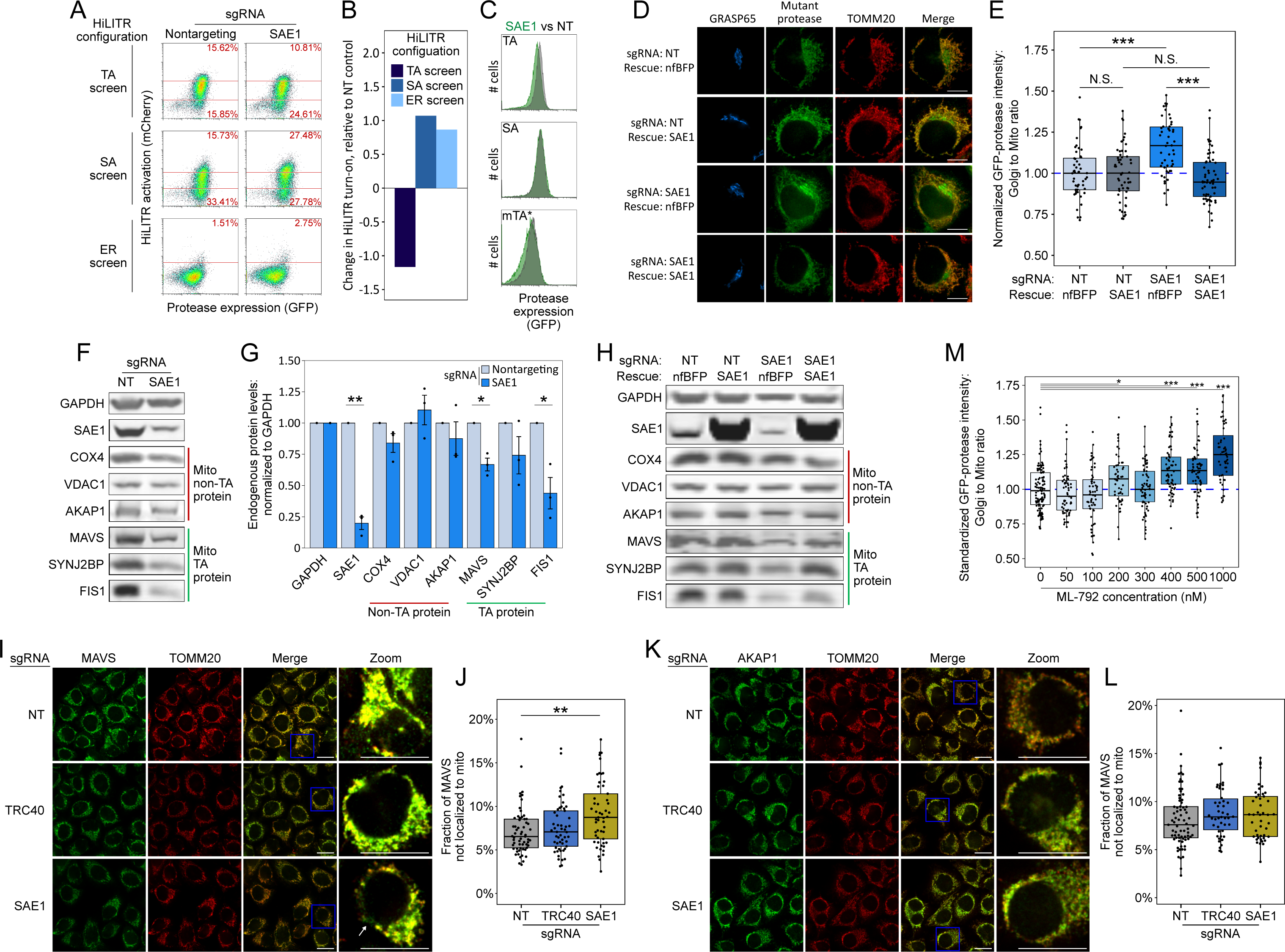
SAE1 knockdown disrupts localization and abundance of mitochondrial tail-anchored proteins. **(A)** SAE1 knockdown by CRISPRi reduces Hi-LITR activation in TA screen configuration, while increasing Hi-LITR activation in SA and ER screen configurations. **(B)** Quantitation of data in (A). **(C)** Expression levels of GFP-tagged mitochondrial proteases from samples in (A). Green, SAE1 knockdown cells; grey, control cells with nontargeting guide. **(D)** SAE1 knockdown increases mislocalization of the GFP-tagged mTA * protease from mitochondria to GoIgi. HeLa cells expressing mTA * protease and dCas9-KRAB were infected with SAE1 sgRNA or NT control for 9 days. In rows 2 and 4, SAE1 knockdown was rescued by overexpression of sgRNA-resistant SAE1. nfBFP, non-fluorescent BFP. Mitochondria and Golgi are visualized with anti-TOMM20 and anti-GRASP65 antibodies, respectively. Scale bar, 10 µm. **(E)** Quantitation of data in (D) along with ∼20 additional fields of view (n = ∼50 cells per condition). N.S. = not significant, ***p < 0.001, Student’s t-test. **(F)** SAE1 knockdown by CRISPRi reduces abundance of some endogenous mitochondrial TA proteins. HeLa cells were infected with SAE1 sgRNA or nontarge t control for 9 days before lysis and Western blot detection of tail-anchored mitochondrial proteins (MAVS, SYNJ2BP, FIS1) and non-tail-anchored mitochondrial proteins (COX4, VDAC1, AKAP1). Uncropped blots in Figure S17. **(G)** Quantification of data in (F) along with two additional biological replicates. Error bars = SEM. *p < 0.05, **p < 0.01, Student’s t-test. **(H)** Overexpression of sgRNA-resistant SAE1 in SAE1 knockdown cells partially restores levels of endogenous tail-anchored mitochondrial proteins MAVS and FIS1. Uncropped blots in Figure S18. **(I)** SAE1 knockdown increases the fraction of the endogenous TA protein MAVS that is not localized to the mitochondria. HeLa cells were infected with nontargeting control or sgRNA against TRC40 or SAE1 for 9 days. Endogenous MAVS and the mitochondrial marker TOMM20 were visualized by immunostaining. Zoom-in images are contrast-enhanced. White arrows point to non-punctate MAVS signal in regions without mitochondrial density. Scale bar, 20 µm. **(J)** Quantitation of data in (I) along with ∼5 additional fields of view (n = ∼60 cells per condition). **p < .01, Student’s t-test. **(K)** SAE1 knockdown does not affect mitochondrial localization of the endogenous signal - anchored protein AKAP1. As in (I), with immunostaining of AKAP1 instead of MAVS. Scale bar, 20 µm. **(L)** Quantitation of data in (K) along with ∼5 additional fields of view (n = ∼60 cells per condition). No results were significant by Wilcoxon rank-sum test. **(M)** Chemical inhibition of SUMOylation increases mislocalization of the GFP-tagged mTA* protease from mitochondria to Golgi. Immunofluorescence images are shown in Figure S 11I. Cells were treated with SUMO E1-ligase inhibitor ML-792 cells for 6 days before expression of mTA* protease for 1 day. *p < 0.05, ***p < 0.001, Student’s t-test.

**Figure 5.**
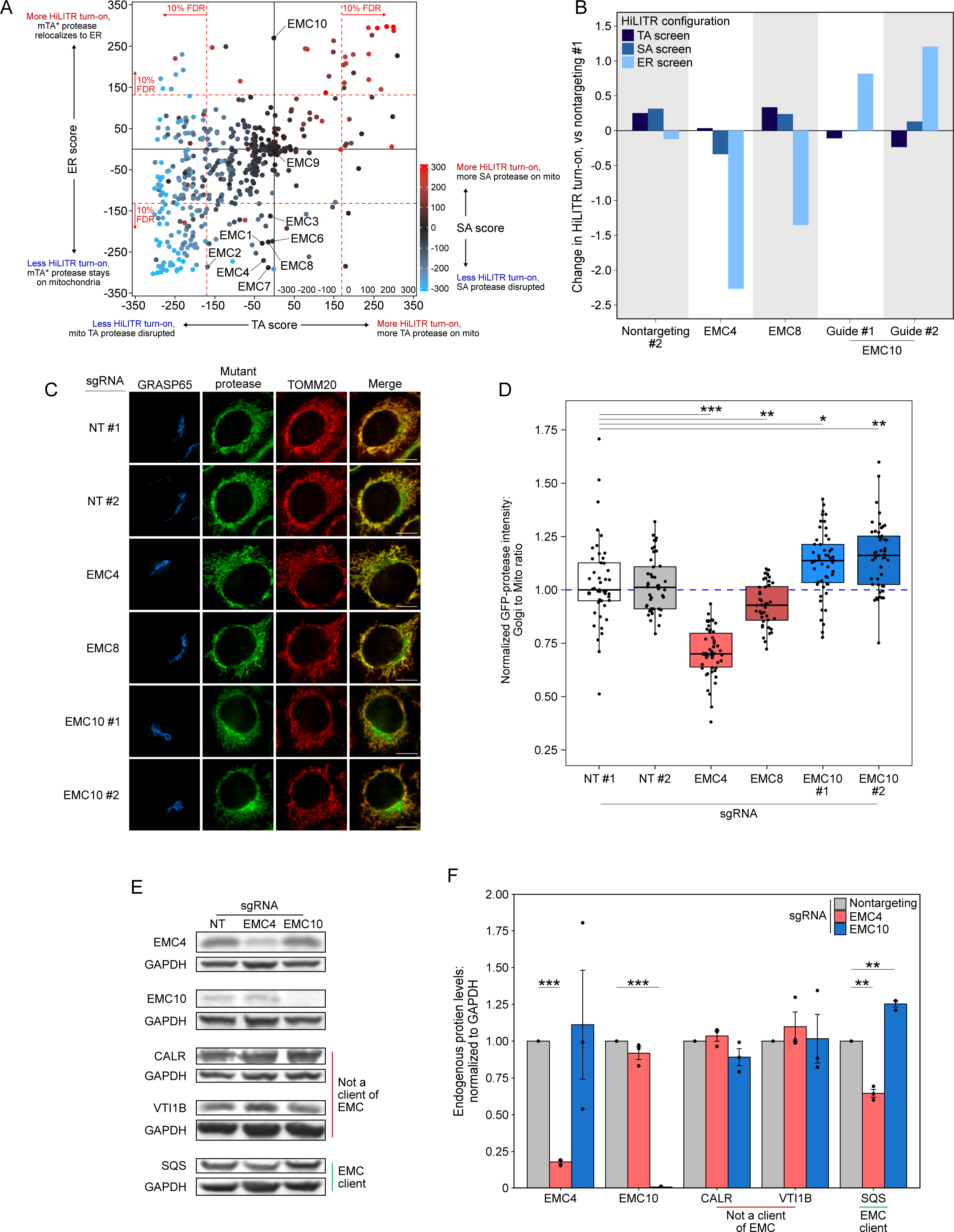
EMC10 has a different regulatory effect on ER-targeted tail-anchored proteins than other EMC subunits. **(A)**Locations of 9 of 10 EMC components in the 3-CRISPRi screen CasTLE plot from Figure 3A. EMC5 was not included in the screen. **(B)** Knockdown of individual EMC subunits (4, 8, and 10) in TA, SA, and ER HiLITR cell lines. FACS data shown in Figure S19A. **(C)** Knockdown of EMC10 increases, while knockdown of EMC4 or EMC8 decreases, the mislocalization of GFP-tagged mTA * protease from mitochondria to Golgi in HeLa cells. Golgi and mitochondria are detected with anti-GRASP65 and anti-TOMM20 antibodies. Scale bar, 10 µm. **(D)** Quantification of data in C along with ∼20 additional fields of view per condition (∼50 cells per sample). *p < 0.05, **p < 0.01, ***p < .001, Student’s t-test. **(E)** Knockdown of EMC subunits has different effects on endogenous EMC client protein SQS. HeLa cells expressing the indicated sgRNAs (EMC10 sgRNA #2) were lysed after 9 days and blotted for the endogenous tail - anchored EMC client protein SQS as well as two non-client proteins (ER lumen protein CALR and ER tail-anchored protein VTI1B). Uncropped blots in Figure S15. **(F)** Quantification of data in (E) along with 2 additional biological replicates per condition. Error bars = SEM. **p < 0.01, ***p < .001, Student’s t-test.

We then searched our data for novel genes that might preferentially influence the targeting of mitochondrial TA proteins. Such proteins could be expected to have a low TA score, because sgRNA-mediated depletion of TA protease from the OMM would reduce HiLITR activation, but medium to high SA score, because SA protease would be minimally affected. A high ER score could also be expected due to a shift in mTA* protease localization from OMM to ERM, resulting in activation of ER-localized TF (Figure 3B). Interestingly, only a single gene met all three criteria: SAE1. SAE1 is an essential protein and a member of the SAE complex (along with SAE2/UBA2), which acts as the sole SUMO E1 ligase in mammalian cells. Modification of target proteins with the small protein tag SUMO can alter protein subcellular localization (Martin et al., 2007; Matunis et al., 1996), regulate protein stability (Desterro et al., 1998; Krumova et al., 2011), and promote cellular stress response (Golebiowski et al., 2009). Several important mitochondrial proteins are SUMOylation targets, including Parkin (Um and Chung, 2006) and Drp1 (Prudent et al., 2015). Intriguingly, several chaperones implicated in the handling of TA proteins, including the ubiquilins (Itakura et al., 2016) and STIP1, are also SUMOylated (Hendriks et al., 2018; Soares et al., 2013).

As each HiLITR configuration is unlikely to be perfectly sensitive, we also looked for genes that met two of the three criteria (Figure 3B). Several genes involved in mitosis and the cytoskeleton were among the hits with low TA score and mid-to-high SA score, including BORA, CCNK, REEP4, MKI67IP, and SKA1, (Fig. 3C). The quadrant with low TA score and high ER score (Fig. 3D) contained only a few hits, one of which was ATP6V1A, a subunit of the vacuolar ATPase, which has an important role in vesicle trafficking (Dettmer et al., 2006). Additional hits and pathways are discussed in Text S10.

To check the robustness of our hits, we individually validated single sgRNAs in our HiLITR cell lines (Figs. S8A/C and S9, Text S10). In addition to HiLITR activation, we also quantified expression level of the GFP-tagged protease, as mistargeting may result in protein degradation (Fig. S8B/D). We selected SAE1 (Fig. 4) and seven additional hits from Figures 3D/E for validation. Four of these hits (SAE1, CCNK, SKA1, and ATP6V1A) gave robust validation (Figure 4 and S8E), and we found by imaging that knockdown of SKA1 or ATP6V1A increased the fraction of GFP-mTA* protease mislocalized to the Golgi (Fig. S8F/G).

### SAE1 knockdown disrupts localization and abundance of mitochondrial tail-anchored proteins

Validation with individual sgRNAs against SAE1 in HiLITR cell lines recapitulated the results of the CRISPRi screens (Figure 4A/B). In addition, we observed that SAE1 knockdown specifically reduced the abundance of GFP-tagged mito TA protease, but not GFP-tagged mito signal-anchored protease (Fig. 4C). This may be because a significant fraction of mito TA protease that fails to target to the OMM is destabilized and degraded. We also imaged the subcellular localization of GFP-tagged mito mTA* protease by fluorescence microscopy, and found that knockdown of SAE1 causes redistribution of this construct away from the mitochondria to ER/Golgi compartments. The effect was rescued by overexpression of sgRNA-resistant SAE1 (Fig. 4D/E, S11D/E).

We also examined the effect of SAE1 knockdown on *endogenous* mitochondrial proteins. We infected HeLa cells expressing dCas9-KRAB with a nontargeting sgRNA or an sgRNA against SAE1. After 9 days of guide expression, we harvested cells and measured the abundance of endogenous mitochondrial proteins, using GAPDH as a loading control because it is known to not be SUMOylated (Huang et al., 2018). Of three mitochondrial tail-anchored proteins tested (MAVS, SYNJ2BP, and FIS1), two showed significant depletion upon knockdown of SAE1 (Fig. 4F/G, S12). In contrast, three non-tail anchored mitochondrial proteins with diverse targeting signals (COX4, VDAC1, and AKAP1) all showed no reduction in protein abundance upon SAE1 knockdown (Fig. 4F/G, S12). Overexpression of sgRNA-resistant SAE1 partially restored levels of the mitochondrial tail-anchored proteins (Fig. 4H, S13).

We then used fluorescence microscopy to examine the localization of endogenous mitochondrial proteins upon SAE1 knockdown. In control cells expressing nontargeting sgRNA, the tail-anchored protein MAVS almost completely localizes to the mitochondrion (Fig. 4I/J), although some non-mitochondrial MAVS appears in punctate structures that may correspond to peroxisomes (Dixit et al., 2010). Knockdown of the ER-specific chaperone TRC40 did not alter the extent of colocalization between MAVS and the mitochondrial marker TOMM20. In contrast, knockdown of SAE1 significantly increased the non-mitochondrial fraction of MAVS (Fig. 4J). In these cells, we observed non-mitochondrial MAVS in fibrous, perinuclear structures (Fig. 4I, zoom) distinct from the small puncta observed in control samples. In contrast to MAVS, the localization of endogenous AKAP1, a signal-anchored mitochondrial protein, was unaffected by knockdown of SAE1 (Fig. 4K/L). Taken together with the western blotting data (Fig. 4G), our results suggest that SAE1 knockdown triggers both degradation of the endogenous mitochondrial TA protein MAVS, and mislocalization of a portion of this protein to non-mitochondrial compartments.

Knockdown of SAE1 might impair mitochondrial TA protein topogenesis through SUMOylation activity or through undetermined nonenzymatic binding interactions. To distinguish between these possibilities, we examined the effects of the small-molecule inhibitor ML-792, which inhibits global SUMOylation with an EC50 of 19 nM (Huang et al., 2018). Inhibition of SUMOylation in K562 cells for just two days reproduced the effects of SAE1 knockdown on HiLITR activation in the TA, SA, and ER screen cell lines (Fig. S11F-H). Furthermore, in HeLa cells expressing the GFP-tagged mito mTA* protease, inhibition of SUMOylation with ML-792 in excess of 400 nM increased mislocalization of GFP-mTA* protease to Golgi (Fig. 4M, S11I).

### EMC10 is an EMC component with a distinct regulatory effect on ER membrane proteins

In our ER screen, the HiLITR TF is localized to the ER membrane, and the mTA* protease is distributed between the OMM and ERM (Figure S4E). Therefore, the ER screen could also identify regulators of ER TA proteins, whose knockdown could decrease colocalization of the mTA* protease and ER-TF, reducing HiLITR activation. Thus, we closely examined hits that gave large CasTLE scores in the ER screen. Several of the highest scoring genes were components of the ER membrane complex (EMC), which mediates proper insertion of both multipass transmembrane proteins (Chitwood et al., 2018; Shurtleff et al., 2018) and a subset of tail-anchored proteins (Guna et al., 2018) into the ER membrane.

Seven of the 9 EMC subunits included in our screen reduced HiLITR activity in the ER screen (Fig. 5A), but had no effect in the TA or SA screens (EMC subunits 1/2/3/4/6/7/8). This suggests that our mTA* protease is a client of EMC, while the mitochondrial TA and SA proteases do not interact with the EMC. Intriguingly, EMC10 was one of the strongest HiLITR-*increasing* hits in the ER screen (Fig. 5A), an effect in stark contrast to the other EMC components. EMC10 is less well-conserved than other EMC proteins (Wideman, 2015), does not cluster with core components in genetic interaction mapping (Jonikas et al., 2009), and is dispensable for complex stability (Volkmar et al., 2019). EMC10 forms contacts with EMC1 and EMC7 in the ER lumen (O’Donnell et al., 2020), and mutations at the EMC1/7 interface strongly increase the level of a reporter based on the canonical tail-anchored EMC client SQS (Miller-Vedam et al., 2020). Therefore, we wondered if EMC10 might antagonize or regulate the activity of the EMC, such that its depletion would increase the insertion of client tail-anchored proteins.

We began by validating the HiLITR screen results of EMC4, EMC8, and EMC10. We chose EMC4 because it is part of the main cavity (along with EMC3/6) that mediates insertion of EMC substrates (Bai et al., 2020; Pleiner et al., 2020), but its depletion does not destabilize the rest of the EMC (Volkmar et al., 2019). Compared to two nontargeting controls, EMC4 and EMC8 knockdown both decreased HiLITR activity specifically in the ER screen, while two guides against EMC10 both increased HiLITR activity (Fig. 5B, S14A).

To further assess if EMC10 knockdown increases the insertion of client proteins, we used fluorescence microscopy to assess the distribution of GFP-tagged mTA* protease in HeLa cells. While knockdown of EMC4 and EMC8 both decreased the colocalization of mTA* protease with the Golgi, the guides against EMC10 increased the presence of mTA* protease at the Golgi. This suggests that EMC10 knockdown increases insertion of mTA* protease at the ERM relative to the OMM.

Although the mTA* protease appears to be a client of the EMC, it is not derived from a canonical ER tail-anchored protein. To query a more representative construct, we generated two new proteases targeted to the ER membrane via fusion to the native ER TA protein SQS or to its transmembrane domain (Fig. S1, S14B). In K562 cells expressing the HiLITR ER-TF, knockdown of EMC10 knockdown increased HiLITR activation with full length SQS-protease (Fig. S14C-F). We next tested the effects of EMC10 knockdown on endogenous SQS. HeLa cells were infected with guides against EMC4 or EMC10, and cells were harvested after 9 days of sgRNA expression. Western blotting confirmed knockdown of EMC4 and EMC10 (Fig. 5E, S15). Neither EMC4 nor EMC10 knockdown altered the levels of the ER-translocon-dependent lumen protein calreticulin (CALR) or the TRC-dependent tail-anchored protein VTI1B. In contrast, EMC4 knockdown significantly decreased the level of SQS, while EMC10 knockdown significantly increased SQS levels (Fig. 5E/F, S15).

## Discussion

With the advent of CRISPR-based gene perturbation and continued improvements to next-generation sequencing, large-scale functional genomics studies have become simpler, faster, and more cost-effective, particularly for pooled-format screens using conventional equipment and reagents. Presently, the biology which can be accessed by pooled-format screens is most limited by our ability to couple a cellular function of interest to a simple, robust readout. Several recent studies have combined pooled cell culturing with automated high-content microscopy, using in-place sequencing (Feldman et al., 2019; Wang et al., 2019), arrayed imaging (Wheeler et al., 2020), or photoinducible reporters (Kanfer et al., 2021; Yan et al., 2021) to identify hits. Such approaches simplify cell culturing and collection, but identifying phenotypic hits still requires time-consuming microscopy and computational analysis. In this work, we have developed an alternative approach to the study of protein localization with pooled screens. HiLITR is a genetically-encoded tool which provides light-gated readout of protein localization, enabling fast, genome-wide screening with high-coverage, while dispensing with the need for specialized equipment or software.

Protein complementation assays (PCAs), such as split GFP (Cabantous et al., 2005), can also be used as readouts in high-throughput assays. Compared to PCAs, HiLITR provides signal amplification, which may improve sensitivity towards weak or rare interactions. The customizable HiLITR reporter is also not limited to fluorescent reporter production.

We envision a wide range of possible HiLITR applications in future studies. Because HiLITR is modular, it can be easily designed for applications at other organelles. The light-gating of HiLITR can also be applied to the time-resolved detection of transient changes in protein localization, such as protein translocation in response to internal or external cues. HiLITR could also be formatted for use in other high-throughput assays, such as screens of chemical libraries.

While powerful, HiLITR also bears certain limitations. Loss-of-activation screens produce false positives that nonspecifically decrease the expression of HiLITR components. Repeated applications of HiLITR by the scientific community may generate lists of recurring false positives, but false-positives can also be filtered through the use of matched counterscreens (e.g. the SA screen vs the TA screen). Gain-of-activation screens are likely to produce fewer false positives, but they must be designed with prior knowledge of how perturbation will alter construct localization. We also recommend cautious interpretation of hits that produce strong growth defects, for which we have observed generally lower reproducibility.

In this study, we combined HiLITR with CRISPRi to identify genes involved in the trafficking of mitochondrial and ER membrane proteins. Using a pooled format, we performed a whole-genome screen followed by three smaller-scale screens, focusing on the identification of genes which specifically perturb the trafficking of mitochondrial tail-anchored proteins. We identified SAE1, an essential member of the mammalian SUMO E1 ligase. Knockdown of SAE1 specifically decreased the abundance of the endogenous mitochondrial tail-anchored proteins MAVS and FIS1. Knockdown of SAE1 also promoted mislocalization of endogenous MAVS. The effects of SAE1 knockdown were phenocopied by chemical inhibiting SUMOylation. While the effect of SAE1 knockdown appears to be specific to mitochondrial tail-anchored proteins, the perturbation may be indirect. Both STIP1 and the ubiquilins show evidence of SUMOylation (Hendriks et al., 2018), and it is possible that SUMOylation of the chaperones which handle tail-anchored proteins may be functionally relevant.

In our ER screen, we observed a novel and unexpected consequence of EMC10 knockdown. While loss of other EMC subunits decreased the ER localization of our mutant tail-anchored protease, EMC10 knockdown produced the opposite effect, which we confirmed by immunofluorescence. We further showed that knockdown of EMC10 increased abundance of the endogenous ER tail-anchored protein SQS. EMC10 localizes to the ER lumen, and structural studies show that EMC10 interacts with both EMC1 and EMC7. Disruption of the EMC1/7 interface increases the level an SQS-based reporter construct, and it has been suggested that permissiveness of the EMC might be regulated, altering levels of SQS insertion as the cell responds to changes in demand for sterol synthesis (Miller-Vedam et al., 2020). We propose that EMC10 may play a role in this regulatory process, perhaps by stabilizing a less permissive conformation of the EMC.

## Methods

### Mammalian cell culture

HEK293T cells (ATCC) were cultured as a monolayer in a 1:1 DMEM/MEM mixture (Corning 10-017; Corning 15-010) supplemented with 10% fetal bovine serum (Avantor 97068-085) and 1% penicillin-streptomycin (Corning 30-002-CI, final concentration 1 U/mL penicillin and 100 µg/mL streptomycin) at 37 °C with 5% CO_2_. K562 cells (ATCC) were cultured in suspension in RPMI 1640 (Corning 15-040) supplemented with 10% fetal bovine serum (Avantor 97068-085), 1% penicillin-streptomycin (Corning 30-002-CI, final concentration 1 U/mL penicillin and 100 µg/mL streptomycin) and 1% GlutaMAX (Gibco 35050061) at 37 °C with 5% CO_2_, while subject to 30 rpm linear shaking. For large-scale screens, K562 cells were cultured in spinner flasks (BELLCO 1965-83005) while subject to magnetic stirring at 60 RPM. HeLa cells (CVCL_1922) were cultured as a monolayer in Roswell Park Memorial Institute 1640 (Corning 15-040) supplemented with 10% fetal bovine serum (Avantor 97068-085), 1% penicillin-streptomycin (Corning 30-002-CI, final concentration 1 U/mL penicillin and 100 µg/mL streptomycin) and 1% GlutaMAX (Gibco 35050061) at 37 °C with 5% CO_2_. For fluorescence microscopy experiments, cells were plated on 7 mm x 7 mm glass coverslips in 48 well plates. The coverslips were pretreated with 50 mg/mL fibronectin (Millipore FC010) in identical culturing medium and conditions for at least four hours, in order to improve cell adherence.

### Lentivirus generation and stable integration of constructs

Lentivirus was generated by transfection of lentiviral vector (1,000 ng) and packaging plasmids pCMV-dR8.91 (900 ng) and pCMV-VSV-G (100 ng) with 12 µL of polyethyleneimine (PEI, 1 mg/mL; Polysciences 24765-1) into HEK293T cells that had been grown to 60-80% confluence in 6-well plates. Total volume of media was 2 mL per transfection. About 48 hours after transfection, the cell medium was harvested in 0.5 mL aliquots and flash-frozen in liquid nitrogen, then stored at -80 °C. Prior to infection, viral aliquots were thawed at 37 °C.

For larger-scale lentivirus generation, lentiviral sgRNA vector libraries (8,000 ng) and packaging plasmids (8,000 ng total, same weight-ratios as above), with 50 uL of PEI, were transfected into HEK293T cells cultured in 15 cm dishes. Total volume of media was 30 mL per transfection. 48 hours after transfection, the cell medium was harvested, and an additional 30 mL of media was added to each plate. 24 hours later, this media was combined with the media from the 48 hour time point and filtered through a 0.45 um syringe filter (Millipore SLHV033RB; BD 309653). One 15 cm dish was transfected for each 35 million K562 cells to be transduced.

To infect K562 cells, 50-250 thousand cells in log-phase growth were combined with one or two viral aliquots and 1.6 µL of polybrene (10 mg/mL; Millipore TR-1003-G) in a total volume of 2 mL in a 24-well plate format. The plates were subjected to centrifugation at 1,000 x g and 33 °C for 2 hours. For large-scale experiments, infections were performed in 6-well plates, and the volume of reagents and number of plates used was scaled up in proportion to the desired number of cells infected. HeLa cells were infected by adding one or two viral aliquots to the media of a 6-well plate when the cells had grown to 15-30% confluency. For both K562 and Hela Cell, selection was initiated 2 days after infection with 0.5 µg/mL puromycin (Sigma P8833), which was increased in concentration to 1 µg/mL over the next two days and supplied for a total of 3-6 days. Some plasmids instead required selection with blasticidin (Corning 30-100-RB, starting with 4 µg/mL and increased to 8 µg/mL over a selection time course of 5-7 days) or with hygromycin (Corning 30-240-CR, starting with 100 µg/mL and increased to 200 µg/mL over a selection time course of 5-7 days), or with geneticin (ThermoFisher #10131035, starting with 50 µg/mL and increased to 100 µg/mL over a selection time course of 5-7 days).

### Generation of clonal cell lines

To generate clonal K562 cell lines, cell lines were first generated with stable integration and selection of all desired constructs. Cell density was estimated using a Countess II FL automated cell counter, and cells were serially diluted and plated in a 48-well plate format at a target density of .2 cells/well. After 1-2 weeks of expansion, clonal cell lines were selected for desired levels of construct expression and HiLITR response, as assessed by flow cytometry analysis.

Clonal HeLa cell lines were generated from cell cultures with stably integrated and selected constructs. After lifting and separating cells with trypsin (Corning 25-053), serial dilutions were plated on 10 cm cell culture dishes. A day after plating, individual clones were identified with an Olympus CKX31 benchtop inverted microscope. After 1-2 weeks of expansion, previously identified colonies were isolated with a cloning cylinder (Millipore TR-1004), lifted with trypsin, and transferred to a 6-well plate for further expansion. Clonal lines were then selected for desired levels of construct expression, as assessed by flow cytometry.

### Immunofluorescence staining and fluorescence microscopy

Cells were incubated with 400 ng/mL doxycycline (Sigma D9891) upon plating if a doxycycline-inducible fluorescent construct was to be Imaged. Roughly 12-16 hours after plating, cells were fixed with 4% paraformaldehyde (RICCA 3180) in phosphate-buffered saline (PBS) for 15 minutes. For immunofluorescence experiments with K562 cells, the plates containing the cells were subjected to centrifugation at 1,000 x g during fixation. After fixation, cells were washed with PBS, then permeabilized with 0.2% triton X-100 (Sigma T9284) in PBS for 10 minutes. After washing again with PBS, cells were incubated with primary antibody for 1 hour in 2% BSA (Fisher BioReagents BP1600) in PBS, then washed with PBS and incubated with secondary, fluorophore-conjugated antibody in 2% BSA in PBS for 30 minutes, followed by a final wash before imaging. During wash steps, media was removed with vacuum aspiration at the lowest possible pressure setting. On occasions where media removal was performed by hand, additional PBS washes were incorporated between steps.

Imaging was performed with a Zeiss Axio Observer.Z1 microscope with a Yokogawa spinning disk confocal head, Cascade IIL:512 camera, a Quad-band notch dichroic mirror (405/488/568/647 nm), and 405 nm, 491 nm, 561 nm, and 640 nm lasers (all 50 mW). Images were captured through a 63x or 100x oil-immersion objective for the following fluorophores: BFP and Alexa Fluor 405 (405 laser excitation, 445/40 emission), EGFP and Alexa Fluor 488 (491 laser excitation, 528/38 emission), mCherry and Alexa Fluor 568 (561 laser excitation, 617/73 emission), and Alexa Fluor 647 (647 laser excitation, 700/75 emission). Differential interference contrast (DIC) images were also obtained. Image acquisition times ranged from 50-250 ms per channel, and images were captured as the average of 2 or 3 such exposures in rapid succession. Image acquisition and processing was carried out with the SlideBook 5.0 software (Intelligent Imaging Innovations, 3i).

Primary antibodies used in imaging include the following: Anti-V5 (Mouse, Invitrogen R960); Anti-TOMM20 (Rabbit, Abcam ab186735); Anti-GRASP65 (Mouse, Santa Cruz sc-374423); Anti-CANX (Rabbit, Thermo Fisher PA5-34754); Anti-PEX14 (Rabbit,

Proteintech 10594-1-AP); Anti-RCN2 (Rabbit, Thermo Fisher PA5-56542); Anti-AKAP1 (Mouse, Santa Cruz sc-135824); Anti-MAVS (Mouse, Santa Cruz sc-166583). Secondary antibodies used in imaging include the following: Anti-mouse Alexa Fluor 488 (Goat, Invitrogen A11029); Anti-mouse Alexa Fluor 568 (Goat, Invitrogen A11031); Anti-mouse Alexa Fluor 647 (Goat, Invitrogen A21236); Anti-rabbit Alexa Fluor 568 (Goat, Invtirogen A11036); Anti-rabbit Alexa Fluor 405 (Goat, Invitrogen A31556). MitoTracker Deep Red FM (Invitrogen M22426) was also used for imaging.

### FACS analysis and sorting

FACS analysis and sorting of K562 cells were carried out with a SONY SH800S cell sorter equipped with four collinear excitation lasers (405 nm, 488 nm, 561 nm, and 638 nm; all 30 mW), using a 100 µm sorting chip. The 638 nm laser was disabled during experiments. Additional fluorescent cell cytometry analysis of K562 cells and HeLa cells was performed using a BioRad ZE5 cell analyzer with four parallel excitation lasers (405 nm - 100 mW, 488 nm – 100 mW, 561 nm - 50 mW, and 640 nm - 100 mW). For experiments using the SONY SH800S, instrumental analysis and data processing were performed using the SONY Cell Sorter Software, versions 2.1.2 and 2.1.5. For experiments with the BioRad ZE5, instrumental analysis was performed using Everest software version 2.3 (BioRad) and data processing was performed using FlowJo version 10.7.1.

The following scatter conditions and fluorophores were measured in FACS experiments (note that for the collinear laser configuration of the SONY instrument, all three active lasers engage in simultaneous excitation): forward scatter (“FSC”; SONY: 488/17 emission; BioRad: 488 excitation, 488/10 emission), back/side scatter (“BSC”/“SSC”; SONY: 488/17 emission; BioRad: 488 excitation, 488/10 emission), BFP (SONY: 450/50 emission; BioRad: 405 excitation, 460/22 emission), EGFP (SONY: 525/50 emission; BioRad: 488 excitation, 509/24 emission), mCherry (SONY: 600/60 emission; BioRad: 561 excitation, 615/24 emission). For first experiments with a given laser configuration, appropriate single-fluorophore compensation controls were included. The corresponding compensation matrix was applied to future experiments using the same laser configuration.

A short series of gates was used to focus sorting and analysis on the desired population of cells. Live and dead cells were first separated by plotting back/side scatter area (BSC-A/SSC-A) against forward scatter (FSC-A), and dead or dying cells were excluded by drawing a gate which omitted cells with a high BSC/SSC:FSC ratio, yielding population P_1_. From P_1_, single cells were separated from cell doublets by plotting forward scatter height (FSC-H) against width (FSC-W) and drawing a gate around the predominant population with lower FSC-W values, yielding population P_2_. For experiments featuring sgRNA constructs with BFP expression indicator (particularly the experiments with sgRNA libraries), population P_2_ was further resolved into population P_3_ by plotting a histogram of BFP values and collecting only cells with high expression of BFP (and sgRNA by proxy, omitting cells lacking sgRNA or in the bottom 10% of the sgRNA-positive peak of the histogram). Finally, for some experiments in which samples consisted of homogenous cell populations (with identical reporter, transcription factor, TEV protease, and sgRNA constructs integrated), population P_2_ or P_3_ was refined to population P4, the population of cells expressing TEV protease (which was fused to EGFP) by plotting FSC-A against EGFP and drawing a gate around the cluster of EGFP-positive cells.

Samples were maintained on ice prior to instrumental analysis. For the large-scale sorting experiments, the acquisition and collection chambers of the SONY SH800S were maintained at 4 °C. Sorted cells were collected in 15 mL conical tubes containing 5 mL of HEPES-buffered RPMI (Sigma R7388) supplemented with 30% fetal bovine serum (FBS; Avantor 97068-085). During sorting, collections tubes which were filled were subjected to centrifugation at 1,000 x g, and the media was removed and replaced with HEPES-buffered RPMI with 10% FBS. After the conclusion of all sorting, collected cells were pooled by sample and sort condition, pelleted again by centrifugation and removal of media, and flash-frozen in liquid nitrogen or immediately subjected to sequencing library preparation.

### HiLITR activation

For small-scale analyses, K562 cells were grown in 6-well plate format (at 50-500 thousand cells/mL), while for large-scale sorting experiments (such as the whole-genome selection), the cells were grown in T125 flasks (at 300-600 thousand cells/mL). Cells were incubated with 400 ng/mL doxycycline to induce expression of the TEV protease component roughly 16-24 hours prior to light stimulation (experiments where conditions differ noted in the text). Upon addition of doxycycline, the plates or flasks were wrapped completely in tin foil, exposing only the vented flask cap, where applicable. Following doxycycline incubation, cells were stimulated with 450 nm blue light from a 28.8 x 28.8 cm^2^, 22W panel (26.5 mW/cm^2^; Yescom YES3110) for a period of 2-8 minutes (depending on the experiment and sample). Plates or flasks were placed directly on top of the panel and agitated by hand once every 1-2 minutes to prevent settling of cells. Light stimulation was carried out at room temperature in a dark room with only red light sources as additional illumination for visual aid. Following light stimulation, cells were returned to tin foil wrapping and replaced in the 37 °C, 5% CO_2_ incubator for 8-16 hours to allow for expression of the reporter construct. After expression, K562 cells were transferred to appropriate tubes for FACS analysis/sorting and placed on ice. For small-scale experiments, cells were sorted in their native RPMI medium. For the large-scale sorts, the cells were collected by centrifugation at 1,000x and the media was removed and replaced with HEPES-buffered RPMI with 10% FBS. During this medium-replacement step, cells were concentrated to a density of 8-12 million cells/mL.

For HiLITR experiments with HeLa cells, light stimulation was carried out in an identical manner, except cells were cultured on 7 mm x 7 mm glass coverslips, they were not agitated during light stimulation, and they were subjected to immunofluorescence staining and fluorescence microscopy after reporter expression.

### Model Selection

A clonal K562 line was generated with a “matched” HiLITR configuration, bearing a tail-anchored, mitochondrial TEV protease component and a signal-anchored, mitochondrial transcription factor component. The clone was selected on the bases of good light/dark sensitivity, typical TEV protease expression levels, and HiLITR activation that was not atypically robust. Two additional K562 cell lines with “mismatched” HiLITR configurations were generated, bearing the same mitochondrial transcription factor and either a tail-anchored, ER TEV protease component or an NES-tagged, cytosolic TEV protease component. One day prior to selection, the density of each cell line was estimated using a Countess II FL automated cell counter. For each mismatched-HiLITR cell line, the matched-HiLITR clonal line was mixed at a 1:20, 1:2, 5:1 and 50:1 ratio of matched to mismatched cells, creating a <u>calibration series</u> over four orders of magnitude (∼200 thousand cells/sample). Additional 1:20 population mixtures (∼2 million cells/sample), as well as unmixed cell lines (∼200 thousand cells/sample), were then subjected to the HiLITR activation protocol (doxycycline-induced TEV protease expression, 3.5-minute light stimulation). Unmixed cell lines were analyzed by FACS (SONY SH800S), and the resulting activation profiles were used to design gates to maximally enrich the matched-HiLITR cell line from the pooled mixture. Just prior to sorting, a “pre-sort” <u>baseline</u> of the mixed cell line sample was set aside from each pooled mixture. Sorting was conducted for about 20 minutes per sample, and about 150,000 cells were <u>collected</u> per sort. Immediately after sorting, RNA was extracted from the post-sort <u>collected</u>, pre-sort <u>baseline</u>, and <u>calibration series</u> populations. RT-qPCR analysis was performed to measure the levels of matched (mitochondrial) TEV protease transcript and mismatched (cytoplasmic or ER) TEV protease transcript in each sample. Three technical replicates were measured per sample, with the following sequencing primers:

-Forward primer (same for all transcripts): 5’-CATGGTGGAATTCGGTTCCACG-3’
-Mitochondrial TEV protease reverse primer: 5’-GGTGAGGGCCTTCCACTACC-3’
-ER TEV protease reverse primer: 5’-GGACTCCACGGTGGTGATTC-3’
-Cytoplasmic TEV protease reverse primer: 5’-CGGCCAGCTCTCCACTACC-3’

A 60 °C annealing temperature was used in the qPCR reaction. For each sample, the ratio of matched protease transcript to mismatched protease transcript was used as a proxy for the ratio of cells from the corresponding populations. Comparison of the transcript ratios from the pre-sort and post-sort samples to the calibration series enabled calculation of the absolute ratio of matched-protease to mismatched-protease cells in each sample, from which the corresponding enrichment of cells bearing the matched protease could be derived.

### RNA extraction

RNA extraction was performed using a RNeasy Plus Mini Kit (Qiagen 74134). Extraction was performed in accordance with the protocol provided in the kit, at a 350 µL scale for pelleted cells.

### RT-qPCR analysis

To convert RNA to single-stranded cDNA for qPCR analysis, 8 µL of extracted RNA was mixed with 1 µL of 10 mM dNTPs and 1 µL of 50 µM random hexamer primer (Invitrogen N8080127). The sample was heated to 65 °C for 5 minutes and then stored on ice for 1 minute. The sample was then added to a mixture of 1 µL SuperScript III RT, 4 µL 5X First-strand Buffer, and 2 µL 0.1 µM DTT (all from Invitrogen 18080044), with 1 µL RiboLock RNAse inhibitor (Thermo Scientific EO0382), and 2µL RNAse-free water. Single stranded DNA was generated by placing the mixture on a thermocycler for 10 minutes at 25 °C, followed by 1 hour at 55 °C and 15 minutes at 40 °C, before holding at 4°C.

To analyze cDNA by qPCR, cDNA was first diluted 25-fold in water. Subsequently, 2 µL of cDNA was mixed with 2.4 µL water, 0.3 µL each of 10 µM forward and reverse primers, and 5 µL of Maxima SYBR Green/ROX qPCR Master Mix (Thermo Scientific K0221). Samples were prepared on ice and arranged in a MicroAmp 48-well reaction place (Applied Biosystems 4375816), which was sealed with MicroAmp optical adhesive film (Applied Biosystems 4375323). Instrumental analysis was performed on a StepOne Real-Time PCR system (Applied Biosystems 436907) using StepOne Software (version 2.2.2). Samples were quantified over 40 cycles of amplification, followed by melt curve analysis for quality control. Count values for each sample were obtained using automatic thresholding performed by the software, and count values were exported to Microsoft Excel for additional analysis.

### Whole-genome selection

The top 5 sgRNA per gene from a genome-wide CRISPRi library (Horlbeck et al., 2016) were used for the genome-wide CRISPRi screen. After generation of lentivirus, the library was infected into the clonal K562 cell line generated for the model selection (mitochondrial transcription factor, tail-anchored mitochondrial TEV protease). Infection was performed in two technical replicates with 280 million K562 cells. Based on FACS analysis of BFP-positive cells, multiplicity of infection was 0.4, for a theoretical coverage of 1100X per library element. We selected for sgRNA incorporation with puromycin, and 36 hours prior to FACS sorting, induced TEV protease expression with doxycycline. Cells were maintained at or above coverage for the culture duration. The cells were transferred to T150 flasks (60 mL) for light stimulation 12 hours prior to sorting, then returned to the spinner flask for reporter expression. Sorting was performed 9 days after infection. About 200 million cells were analyzed by FACS for each technical replicate. Gates were set to collect cells with the top 15% and bottom 15% of mCherry reporter expression, representing the cells with the greatest and least HiLITR activity. Based on sorting purity parameters, about 18 million cells were collected for each gate. Genomic DNA was harvested from cells (QIAGEN 51192) immediately after completion of sorting.

### Matched sublibrary selections

The library for the three matched selections was designed based on the results from the whole-genome selection. The library featured sgRNAs targeting 586 genes (5 sgRNAs per gene) as well as 500 non-targeting controls. Genes targeted were selected based on significance in the whole genome selection and on lack of clear annotation related to transcription or translation. A smaller set of genes corresponding to known protein trafficking pathways was also included.

In addition to the clonal K562 cell line that was previously used in the whole-genome selection (TA cell line), two additional clonal K562 cell lines were generated. The first clonal cell line expressed the same mitochondrial transcription factor as the whole-genome selection cell line, but the TEV protease was a signal-anchored mitochondrial construct (SA cell line). The second clonal cell line expressed a mutagenized tail-anchored mitochondrial TEV protease which was sensitized for misincorporation into the ER membrane, and the transcription factor construct in this cell line was localized to the ER membrane (ER cell line).

Each cell line was transduced with lentivirus of the sgRNA sublibrary at 50 million cell scale for the TA and SA cell lines, and 70 million cell scale for the ER cell line. Cell lines were split into two technical replicates immediately after infection, before cells began to divide. Multiplicity of infection was estimated as follows, based on proportion of BFP-positive, sgRNA-expressing cells: 0.8 for the TA cell line (5,700 x coverage per technical replicate), 1.4 for the SA cell line (10,000 x coverage), and 0.16 for the ER cell line (1,600 x coverage). Cells were cultured in T150 flasks with linear shaking, and coverage levels were maintained during cell culturing and selection. TEV protease expression was induced with doxycycline 36 hours prior to sorting, and light stimulation was performed 12 hours prior to sorting. Sorting was performed 11 days after infection. For the TA cell line, cells with the top and bottom 16% of mCherry reporter expression were collected (11 million cells per condition). For the SA cell line, cells with the top 16% and bottom 33% of mCherry reporter expression were collected (12 million and 24 million cells, respectively). For the ER cell line, cells with the top 9% and bottom 30% of mCherry reporter expression were collected (6 million and 22 million cells, respectively). Genomic DNA was harvested from cells immediately after completion of sorting.

### Sequencing library preparation

The integrated sgRNA library was PCR amplified and separately barcoded for each collected population with Herculase II Fusion DNA Polymerase (Agilent 600677). Samples were then pooled and sequenced on an Illumina NextSeq flow cell with aligned read counts as follows for each screen:

Whole-genome screen: TR1 (HiLITR active) – 51 million; TR1 (HiLITR inactive) – 51 million; TR2 (HiLITR active) – 49 million; TR2 (HiLITR inactive) – 47 million.

TA screen: TR1 (HiLITR active) – 6 million; TR1 (HiLITR inactive) – 8 million; TR2 (HiLITR active) – 7 million; TR2 (HiLITR inactive) – 8 million

SA screen: TR1 (HiLITR active) – 9 million; TR1 (HiLITR inactive) – 6 million; TR2 (HiLITR active) – 7 million; TR2 (HiLITR inactive) – 9 million

ER screen: TR1 (HiLITR active) – 11 million; TR1 (HiLITR inactive) – 13 million; TR2 (HiLITR active (11 million); TR2 (HiLITR inactive) – 11 million

### Data analysis of CRISPR screens

CRISPR screens were analyzed using CasTLE (Morgens et al., 2016), a maximum likelihood estimator that determines each gene’s effect size based on the enrichment of its sgRNAs relative to a null effect model derived from the enrichments of non-targeting control sgRNAs. The significance of each gene’s effect size is tested by evaluating it against the distribution of the estimated effect sizes from random permutations drawn from all targeting sgRNA within the library.

More explicitly, the effect size is the log_2_-transformed maximum likelihood estimate (using a Bayesian framework) of the change in the ratio of high mCherry to low mCherry cells for knockdown of a given gene, relative to the complement of non-targeting controls. The CasTLE score is twice the log-likelihood ratio of estimated effect size (monotonically increasing with decreasing p-value).

### Cloning individual sgRNA lentiviral vectors

Single-sgRNA vectors were cloned as follows. The U6:sgRNA_Puro-T2A-BFP lentiviral vector was digested with BlpI and BstXI. Primers corresponding to the sgRNA target (F – 5’-TTG-[*Guide*]-GTTTAAGAGC-3’; R – 5’-TTAGCTCTTAAAC-[*Guide-rev.comp.*]-CAACAAG-3’) were mixed (10 uL of 10 µM oligo each) and annealed by heating to 95°C for 5 min, followed by cooling to 25 °C at 5 °C/minute. Annealed primers were then diluted 1:20 in water and cloned into the digested vector by T4 ligation. Double-sgRNA vectors were cloned as follows. First, a single-sgRNA vector was cloned with the desired target (sgRNA #1), then digested with XhoI and BamHI. Next, the U6:sgRNA_G418 vector was digested with BbsI. Primers corresponding to sgRNA target (F – 5’-ACCG-[*Guide*]-3’; R – 5’-AAAC-[*Guide-rev.comp.*]-3’) were mixed, annealed, and ligated into the vector (“sgRNA #2”). Finally, sgRNA #2 and its corresponding promoter were digested out from the vector with XhoI and BamHI and ligated into the sgRNA #1 vector by T4 ligation.

See Table S1 for specific *guide* sequences used.

### mTA* protease immunofluorescence quantification

A clonal HeLa cell line was generated, expressing dCas9-KRAB-BFP and the doxycycline-inducible, mutant tail-anchored TEV protease fused to EGFP. The clonal line was separately infected with sgRNAs against genes of interest and nontargeting control. After selection of sgRNA-positive cells with puromycin, samples were plated on coverslips on the eighth day after infection, and TEV protease expression was induced with doxycycline. The following day, cells were fixed, permeabilized, and subjected to immunostaining. Primary: mouse anti-GRASP65 (Golgi; Santa Cruz sc-374423, 1:500 dilution) and rabbit anti-TOMM20 (Mitochondria; Abcam ab186735, 1:500 dilution); secondary: goat anti-mouse Alexa Fluor 647 (Invitrogen A21236, 1:1000 dilution) and goat anti-rabbit Alexa Fluor 568 (Invitrogen A11036, 1:1000 dilution).

The resulting images were analyzed with SlideBook 5.0 software (Intelligent Imaging Innovations, 3i), as follows. Cells which were completely present in the image and which had clear signal in the fluorescent channels corresponding to GRASP65, TOMM20, and the mutant tail-anchored TEV protease were bounded to generate unique image objects. For each object, a mask was created on pixels which exceeded a threshold GRASP65 signal. A second mask was created for pixels which exceeded a threshold TOMM20 signal but not the GRASP65 threshold. Average EGFP intensity was calculated for each mask. After measuring all cells, the ratio of average EGFP intensity between the two masks was determined for each cell, then normalized to the median value for cells bearing the non-targeting control sgRNA. Significance was measured by comparing sample distributions to the non-targeting control using an independent two-sample t-test with equal variance assumption.

In some figures, images were taken across multiple experiments and data is combined in the final figure. This is noted where applicable. Data was combined only for presentational purposes, and not in a manner that affected measurements or statistical calculations. In all cases, all images corresponding to an individual guide were obtained in the same experiment, and each experiment had a corresponding nontargeting control sample. Prior to combination of the experimental data, data within each experiment was standardized to a mean of 1 and standard deviation of 0.16 for the nontargeting control sample. Significance for samples in combined-data figures was calculated with respect only to the data from the nontargeting control sample from the same experiment, not the combined sample.

When analyzing localization of the 6 candidate mutant tail-anchored proteases, a Golgi marker was not present. Instead, the Pearson coefficient between the protease channel and the TOMM20 channel was calculated for each cell.

### Endogenous protein immunofluorescence quantification

HeLa cells expressing dCas9-KRAB-BFP were infected with sgRNAs and passaged for 8 days with puromycin selection, before plating on glass coverslips. The following day, cells were fixed, permeabilized, and immunostained. Primary: mouse anti-MAVS (Santa Cruz sc-166583, 1:200 dilution) or mouse anti-AKAP1 (Santa Cruz sc-135824, 1:200 dilution) and rabbit anti-TOMM20 (Abcam ab186735, 1:500 dilution). Secondary: goat anti-mouse Alexa Fluor 488 (Invitrogen A11029, 1:1000 dilution) and goat anti-rabbit Alexa Fluor 568 (Invitrogen A11036, 1:1000 dilution).

The resulting images were analyzed with SlideBook 5.0 software (Intelligent Imaging Innovations, 3i), as follows. Cells which were completely present in the image and which had clear signal in the fluorescent channels corresponding to MAVS and TOMM20 were bounded to generate unique image objects. For each object, a mask was created for pixels which exceeded a threshold intensity for both MAVS and TOMM20, based on intensity in the nucleus. A second mask was created for pixels which exceeded the threshold for MAVS but not TOMM20. After background subtraction (using a non-cellular region), the percent of non-mitochondrial MAVS was measured as the total MAVS intensity from the MAVS-only mask divided by the sum total MAVS intensity across both masks. Significance was measured by Wilcoxon rank-sum test due to the presence of outliers and skew with certain sgRNAs. AKAP1-stained cells were analyzed in the same manner as MAVS-stained cells.

### Western blots

HeLa cells expressing dCas9-KRAB-BFP were infected with sgRNAs and passaged for 9 days in T25 or T75 flasks with puromycin selection. To harvest, cells were washed twice with DBPS. In 2 mL of DPBS, the cells were dislodged with a cell scraper (ThermoFisher 179693) and pelleted by centrifugation (500x g for 3 minutes). The cell pellets were then resuspended in 1 mL DPBS and pelleted again by centrifugation in 1.5 mL Eppendorf tubes (500x g for 3 minutes). Supernatant was removed by aspiration, and the pellets were flash frozen with liquid nitrogen and stored at -80°C. Later, the pellets were lysed by resuspending in RIPA buffer (50 mM Tris pH 8, 150 mM NaCl, 0.1% SDS, 0.5% sodium deoxycholate, 1% Triton X-100 (Sigma T9284) in the presence of 1× protease inhibitor cocktail (Sigma-Aldrich P8849) and 1 mM PMSF. The Eppendorf tubes were incubated for 15 min at 4 °C and vortexed every 3 min for proper sample digestion. Lysates were clarified by centrifugation at 10,000 r.p.m. for 15 min at 4 °C. Protein Loading buffer (6x, 20 uL) was mixed with 100μL of the clarified lysate and boiled for 3 min prior to PAGE gel separation.

Proteins were separated on 9% or 12% SDS-PAGE gels in Tris-Glycine buffer and then were transferred into PVDF membrane (Sigma 05317). The blots were then blocked in 3% BSA (w/v) in TBS-T (Tris-buffered saline, 0.1% Tween 20) for 45 min at room temperature. Blots were then incubated with primary antibody in 3% BSA (w/v) in TBS-T for 1 at room temperature, washed two times with TBS-T for 10 min each, then stained with secondary antibody in 3% BSA (w/v) in TBS-T for 45 min at room temperature. The blots were washed four times with TBS-T for 5 min each time before imaging on Licor Odyssey CLx imaging system. Quantitation was performed using the software provided by Licor.

#### Primary

-SAE1 knockdowns: Mouse anti-GAPDH (Santa Cruz sc-32233) – 1:3000; Rabbit anti-SAE1 (Sigma SAB4500028) – 1:500; Rabbit anti-COX4 (Abcam ab16056) – 1:1000; Mouse anti-VDAC1 (Abcam ab14734) – 1:500); Mouse anti-AKAP1 (Santa Cruz sc-135824) – 1:500; Mouse anti-MAVS (Santa Cruz sc-166583) – 1:250; Rabbit anti-SYNJ2BP (Sigma HPA000866) – 1:500; Rabbit anti-FIS1 (Thermo Fisher 10956-1-AP) – 1:1000)

-EMC knockdowns: Mouse anti-GAPDH (Santa Cruz sc-32233) – 1:4500; Rabbit anti-CALR (Thermo Fisher PA3900) – 1:500; Rabbit anti-VTI1B (Abcam ab184170) – 1:250; Rabbit anti-SQS (Abcam ab195046) – 1:500.

#### Secondary

-SAE1 knockdowns: Goat anti-Mouse IgG IRDye 680RD Polyclonal Antibody (Licor 926-68070) and Goat anti-Rabbit IgG IRDye 800CW Polyclonal Antibody (Licor 926-32211) – 1:20,000

-EMC knockdowns: Goat anti-Mouse IgG IRDye 800CW Polyclonal Antibody (Licor 926-32210) and Goat anti-Rabbit IgG IRDye 680RD Polyclonal Antibody (Licor 926-68071) – 1:20,000

### Resource Availability

Lead contact: Further information and requests for resources or reagents should be directed to the lead contact, Alice Ting

Materials availability: Plasmids generated in the study have been deposited to Addgene or are available upon request (Table S1)

Data and code availability: Available upon request

## Acknowledgments

We thank Tina Kim and Kelvin Cho for comments and feedback during manuscript preparation, Wenjing Wang for technical guidance, Shuo Han for advice on clonal cell line generation, and Kaitlyn Spees for assistance with cloning sgRNA library cloning. We are grateful to Chan Zuckerberg Biohub Stanford for access to FACS sorters. This work was supported by NIH R01 MH119353 (A.Y.T.), NIH Director’s New Innovator Award 1DP2HD084069-01 (M.C.B.), NSF GRFP DGE 1656518 (D.Y.), the Stanford Bio-X Graduate Fellowship Program (R.C.), the NIST JIMB training program (R.C.), NIH 2T32HG000044 (R.C. & D.Y.), a Stanford Center for Systems Biology seed grant (R.C. & D.Y.), and an EMBO long-term postdoctoral fellowship ALTF 1022-2015 (M.I.S.).

## Author Contributions

R.C., D.Y., M.C.B., and A.Y.T. conceived the project. R.C., D.Y., J.S.W., M.C.B., and A.Y.T. designed experiments. R.C. and D.Y. performed all experiments, unless otherwise noted. M.I.S. assisted in immunofluorescence and western blotting experiments. E.T.S. assisted in HiLITR and immunofluorescence experiments. R.C., D.Y., M.C.B., and A.Y.T. interpreted data, and wrote the manuscript with edits from all authors.

---

**Figure S1.**
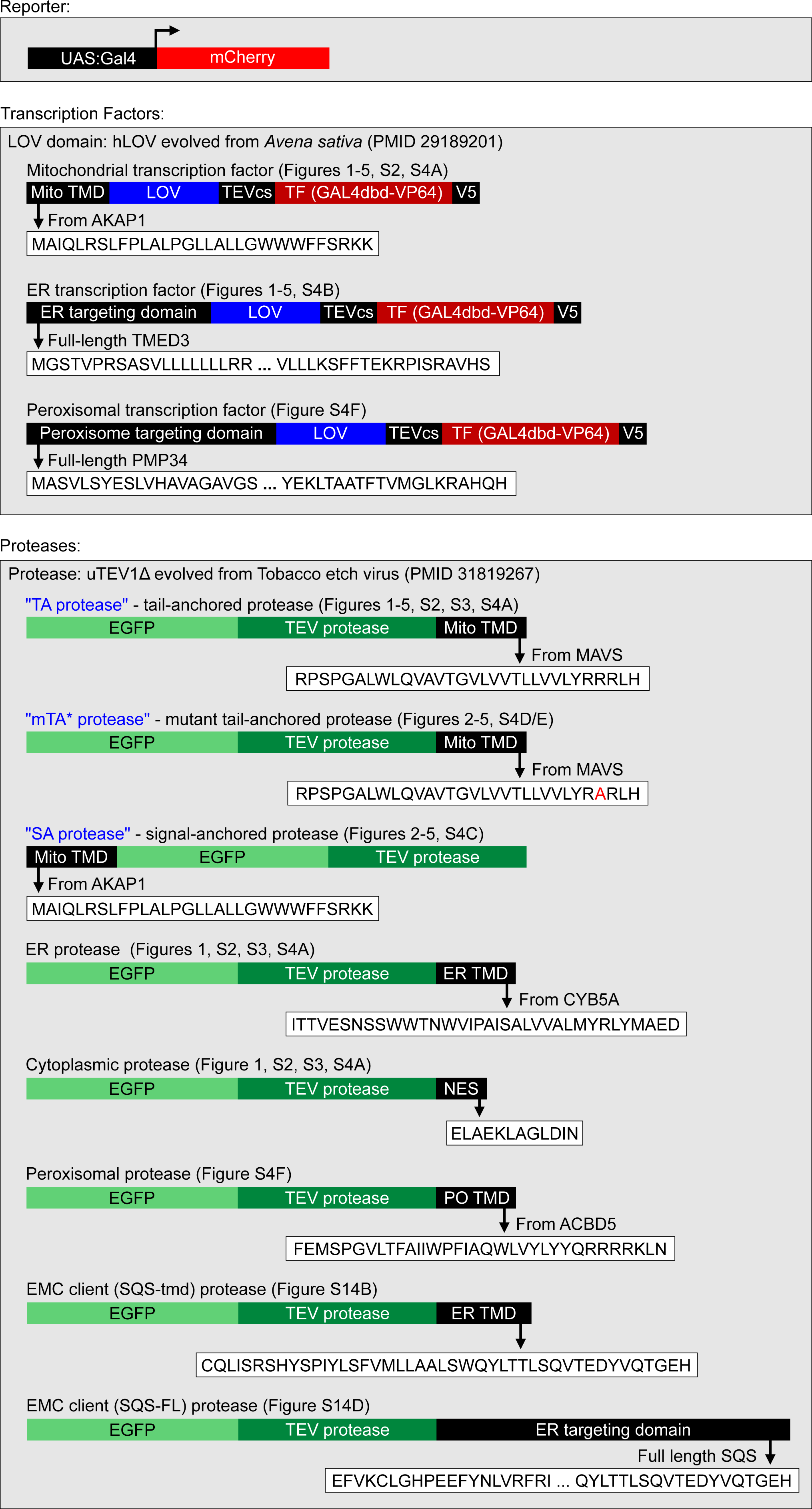
Details for HiLITR constructs used in this study. LOV, TF, and TEV protease domains used in figures S2/3 (HiLITR optimization) vary slightly. Any differences are shown and discussed in the text.

**Figure S2.**
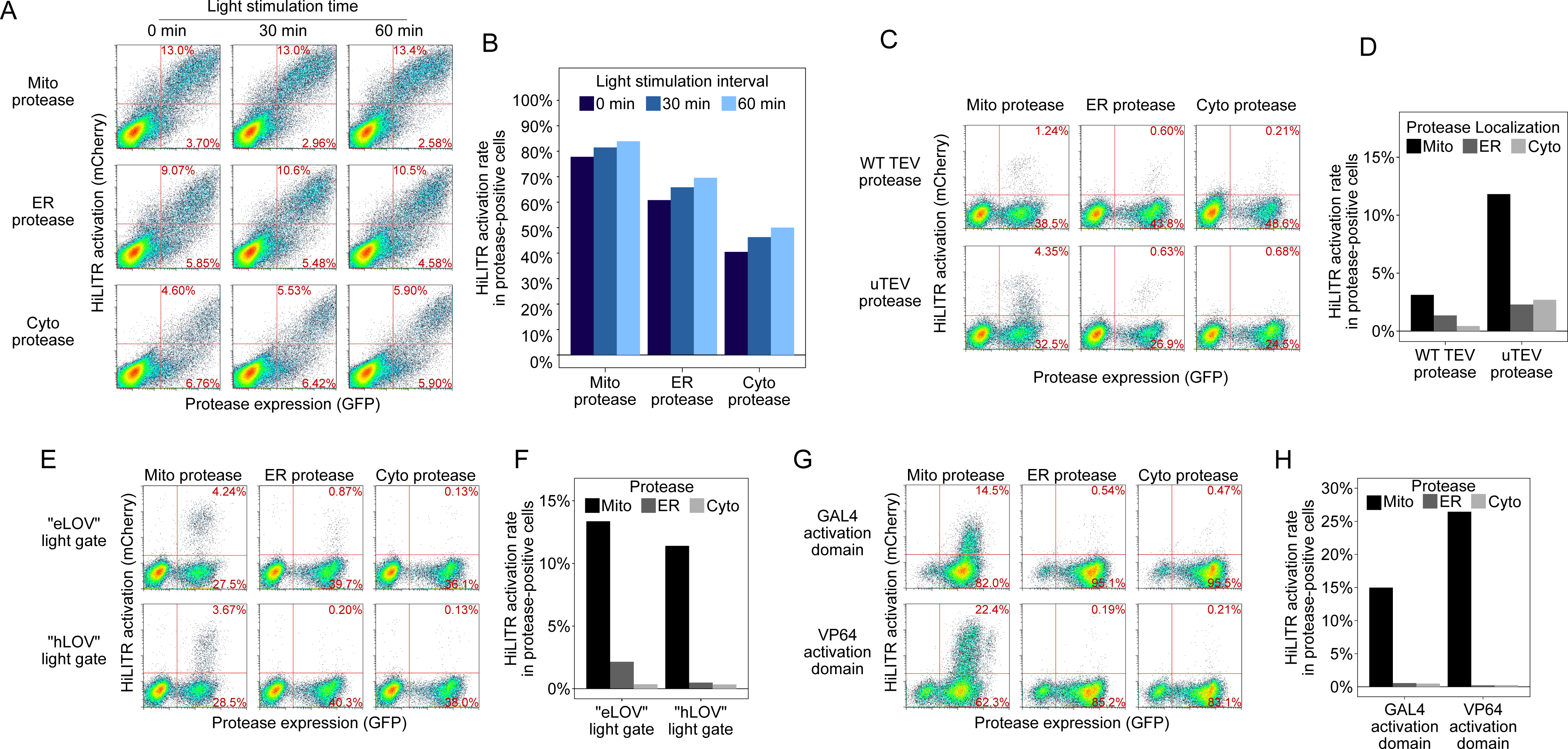
Optimization of HiLITR components. A. FACS plots of HEK cells transfected with HiLITR components. TF is on the outer mitochondrial membrane (OMM), while protease is localized to the OMM (top row), ER membrane (middle), or cytosol (bottom). mCherry on the y-axis reports HiLITR turn-on, while GFP on the x-axis reports protease expression level. In (A), (C), (E), and (G), the percentage of cells expressing protease with or without subsequent reporter expression is shown in the top right and bottom right quadrant of each plot, respectively. TF configuration: “eLOV” (Wang et al., 2017) light gate and GAL4 activation domain. Protease configuration: truncated TEV protease (Wang et al., 2017). B. Quantitation of the results in (A). The fraction of cells expressing both protease and reporter was divided by the total fraction of protease-expressing cells. In (D), (F), and (H), quantitation is performed in the same way. C. FACS plots of K562 cells stably expressing HiLITR TF and mCherry reporter. Cells were transduced with mitochondrial, ER, or cytosolic protease. Truncated TEV protease (top row) and uTEV1Δ protease (Sanchez and Ting, 2020) bottom row) were used. 2 minutes of light stimulation. D. Quantitation of the results in (C). E. FACS plots of K562 cells stably expressing mCherry reporter and “eLOV” or “hLOV” (Kim et al., 2017) TF light gate variant. Cells were transduced with evolved uTEV protease targeted to the mitochondria, ER, or cytosol. No light stimulation was used. F. Quantitation of the results in (E). G. FACS plots of K562 cells stably expressing mCherry reporter, protease, and GAL4 or VP64 TF activation domain variant. No light stimulation was used. H. Quantitation of the results in (G).

**Figure S3.**
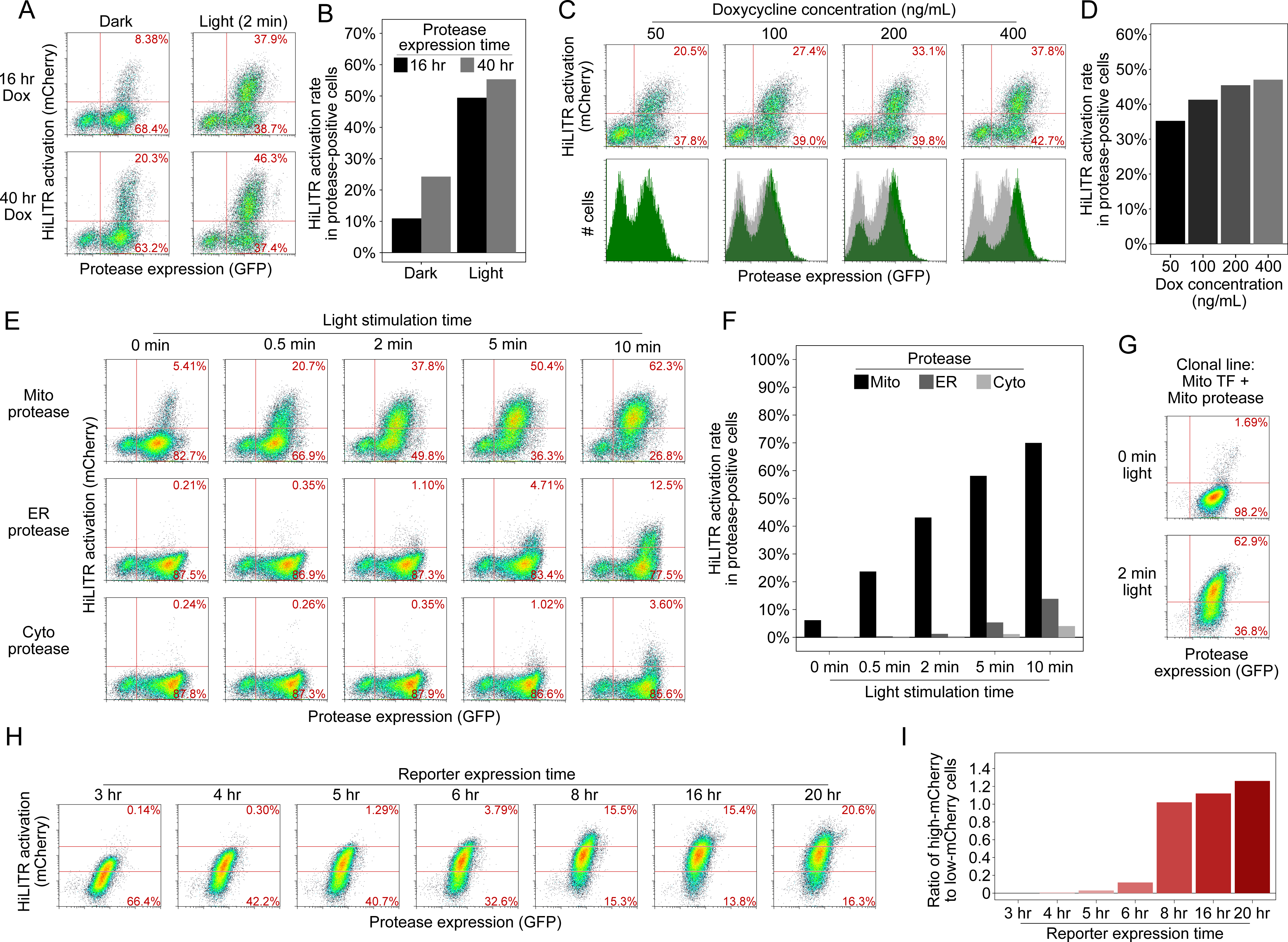
Optimization of HiLITR experimental parameters. A. FACS plots of K562 cells stably expressing mitochondrial protease and HiLITR TF. Protease expression was induced with doxycycline for either 16 hours or 40 hours, and cells were provided with either no light stimulation or two minutes of light stimulation. In (A), (C), (E), and (H), the percentage of cells expressing protease with or without subsequent reporter expression is shown in the top right and bottom right quadrant of each plot, respectively. B. Quantitation of the results in (A). The fraction of cells expressing both protease and reporter was divided by the total fraction of protease-expressing cells. In (D), (F), and (I), quantitation is performed in the same way. C. FACS plots of K562 cells stably expressing mitochondrial HiLITR components. Protease expression was induced for 16 hours with 50-400 ng/mL doxycycline. Light stimulation was provided for 2 minutes. HiLITR activation (top row) and total protease expression (bottom row, with 50 ng/mL doxycycline condition overlayed in gray) were measured across conditions. D. Quantitation of the results in (C). E. FACS plots of K562 cells stably expressing mitochondrial HiLITR components. Light stimulation was varied between 0 and 10 minutes. F. Quantitation of the results in (E). G. FACS plots of a clonal K562 cell line stably expressing mitochondrial TF, mitochondrial protease, and mCherry reporter (compare to Mito protease, top row of (E)). H. The clonal cell line in (H) was stimulated with light for 3 minutes, followed by a variable interval of reporter expression time prior to FACS analysis. The percentage of cells with high mCherry expression (above the top red line) or low mCherry expression (below the bottom red line) is shown in each plot. I. Quantitation of the results in (G).

## Text S3: Optimization of HiILTR experimental parameters Related to Figure S3

After optimization of HiLITR components to minimize background and optimize dynamic range, we investigated the modulation of experimental parameters in the HiLITR assay. First, we looked at expression of the protease. The HiLITR protease is under the expression of a doxycycline inducible promoter to avoid prolonged stable expression of both HiLITR components and to enable cell culturing in ambient light prior to induction of the protease. Reducing the protease expression time window from 40 hours to 16 hours prior to light stimulation improved the signal to noise ratio between the light and dark states from 2.3-fold to 4.9-fold with 2 minutes of light stimulation, with only ∼10% reduction in activation in the light state (Fig. S3A/B). Varying the concentration of doxycycline used to induce protease expression had a modest impact on HiLITR activation, the proportion of protease-positive cells, and total protease expression (Fig. S3C/D).

Next, we asked how HiLITR performance varied with light stimulation time. By varying light stimulation time from 0 to 10 minutes, we found that we could achieve robust HiLITR activation with the mitochondrial protease while maintaining low background with the ER and cytoplasmic proteases with just 2-5 minutes of light stimulation time (Fig. S3E/F). In this experiment, 2 minutes of light stimulation gave a +/-light signal to noise ratio of 7x and a +/-colocalization signal to noise ratio of 35x (vs. ER protease). To improve light vs dark signal to noise, we considered that in the heterogenous population of cells, there were likely some cells which produced light-independent cleavage and other cells which never produced TF cleavage under even extended light stimulation. Reducing cell-to-cell variability is desirable in gene-perturbation studies, so we tested if we could produce a clonal cell line with improved light vs dark signal to noise. Indeed, after generating cell lines, we identified a clone which gave only 1.7% activation in the dark state but 63% activation with two minutes of light stimulation (Fig. S3G), a signal to noise ratio of 37x. Because this clone showed lower activation in the dark state and higher activation in the light state than the heterogeneous population, we reasoned that it must represent an intermediate level of HiLITR sensitivity.

Finally, we tested the change in HiLITR readout with respect to time of reporter expression after light stimulation. In large screens, time of FACS sorting is non-negligible, so it is important to have a readout that is stable with time. With our clonal line, we found that a minimum of 8 hours is required for robust reporter expression, and reporter levels are stable between 8-20 hours post-stimulation (Fig. S3H/I). It is likely that keeping cell samples on ice after 8 hours of reporter expression further stabilized total reporter levels in our high-throughput screens.

**Figure S4.**
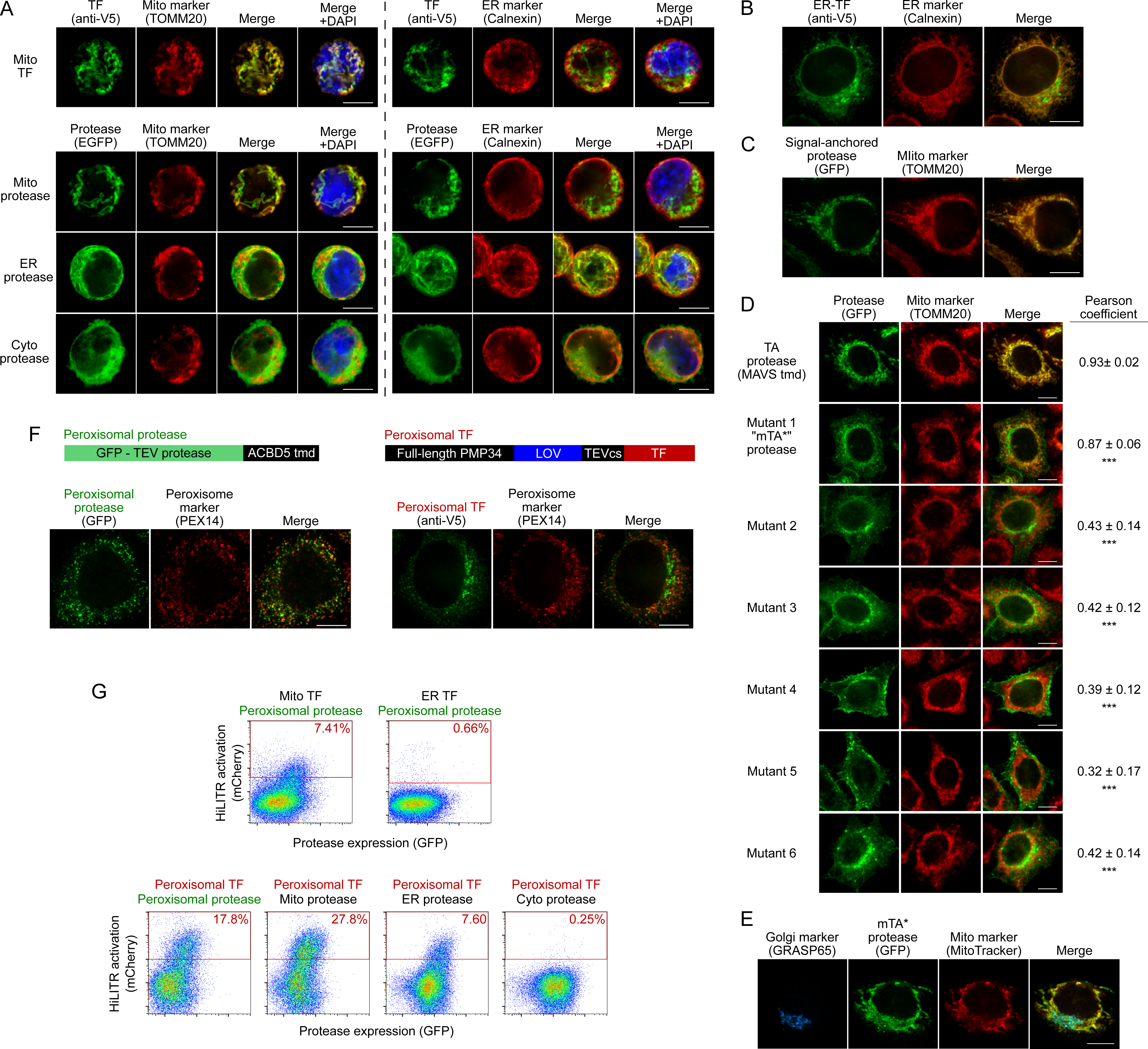
Characterization of HiLITR constructs and cell lines. A. Immunofluorescence microscopy of HiLITR components used in FACS experiment in Figure 1D. The localizations of the mitochondrial TF (V5 tag, top row) and protease constructs (bottom 3 rows) were compared to nucleus (DAPI), mitochondria (TOMM20) and ER (Calnexin) in K562 cells. Scale bar, 10 µm. B. Immunofluorescence microscopy of the ER-localized HiLITR TF used in Figures 1D, 2D, and 2E (“ER transcription factor” in Figure S1). In HeLa cells, the localization of the ER transcription factor (V5 tag) was compared to an ER marker (Calnexin). A fraction of the ER-TF localizes to a non-ER region, consistent with the dual localization of Tmed3 (from which the targeting domain was derived) to ER and Golgi membranes (Emery et al., 2000; Jenne et al., 2002). Scale bar, 10 µm. C. Immunofluorescence microscopy of the signal-anchored mitochondrial protease used in Figures 2C and 2E (“Signal-anchored protease” in Figure S1). In HeLa cells, the localization of the signal-anchored protease (GFP) was compared to a mitochondrial marker (TOMM20). Scale bar, 10 µm. D. Immunofluorescence microscopy of the mutant mitochondrial tail-anchored protease (Mutant 1, “mTA* protease” in Figure S1) and variants (Mutants 2-6; sequences in Methods). HeLa were stained with anti-TOMM20 to visualize mitochondria. Scale bars, 10 µm. At right, mean and standard deviation for Pearson correlation coefficient between the protease and mito marker channels (n = 10-30 cells per condition). **p < .01, ***p < .001, vs TA protease, Wilcoxon rank-sum test. E. Same as D (“mTA* protease”, Mutant 1) but with additional Golgi stain (anti-GRASP65). Scale bar, 10 µm. F. HiLITR constructs for detection of protein colocalization at the peroxisome. Top: domain structures of peroxisome-targeted HiLITR TF and protease constructs. The TF and protease domains face the cytosol. Bottom: Localization of HiLITR constructs in HeLa, using PEX14 peroxisomal marker. Scale bars, 10 µm. G. FACS analysis of K562 cells expressing the indicated HiLITR combinations, 8 hours after 3-minute light stimulation. Percentage of cells in the red gate is quantified in each plot. Mito-TF and ER-TF data was obtained as part of the experiment in Figure 1D.

## Text S4A. HiLITR at the peroxisomal membrane. Related to Figures S4F/G

As part of our HiLITR panel in Figure 1D, also tested the mitochondrial TF and ER-TF against a peroxisomal protease (localization in Fig. S4F). As expected, there was no HiLITR activation with the ER-TF by the peroxisiomal protease (Fig. S4G). Interestingly, the peroxisomal protease did induce mild HiLITR activation with the mitochondrial TF (Fig. S4G). This may be a function of the prevalence of mitochondria-peroxisome contact sites (Chen et al., 2020), which could produce cross-talk between protease and TF constructs on neighboring membranes.

We also attempted to generate a HiLITR TF for the peroxisomal membrane. Unfortunately, despite designing several distinct constructs (including both minimal targeting domains and full-length protein fusions), even the most faithfully targeted construct displayed obvious and profound mistargeting to non-peroxisomal compartments (Fig. S4F). Peroxisomes are formed in a process that involves both the ER and the mitochondrial membrane (Sugiura et al., 2017), and many peroxisome membrane proteins insert at one or both locations, to be subsequently trafficked into newly-derived peroxisomes. It is likely that overexpression of the peroxisomal TF construct produces pools of TF on the mitochondria and/or ER that are too abundant to be efficiently concentrated into nascent peroxisomal membranes. The phenomenon of peroxisomal fusion constructs mislocalizing to the mitochondria or ER has been previously observed (Kim et al., 2006; Sugiura et al., 2017). Consistent with the incomplete targeting of the peroxisomal TF to the peroxisome, we observed HiLITR activation when paired with the peroxisomal, mitochondrial, or ER proteases (Fig. S4G). Importantly, cytoplasmic protease did not activate reporter expression with the peroxisomal TF (Fig. S4G), indicating that colocalization is still a requirement for TF release.

## Text S4B. Generation of the mutant tail-anchored mitochondrial protease (mTA* protease)

For the ER screen (Figure 2D), we sought to generate a mutant tail-anchored mitochondrial protease with a propensity to mistarget to the ER. To do this, we considered the features of tail-anchored proteins which promote ER vs mitochondrial targeting. Tail-anchor sequences of native ER proteins tend to be longer, more hydrophobic, and have fewer basic flanking residues than mitochondrial tail-anchor sequences (Beilharz et al., 2003; Costello et al., 2017; Horie et al., 2002). We found that neutralizing just one of three positive residues flanking the transmembrane domain in our MAVS-based mito TA protease produced detectable mislocalization to the ER and Golgi (Mutant 1, Fig. S4D-E), while other mutations disrupted mitochondrial localization too severely (Fig. S4D, Table S1). Note that localization of protease variants in S4D to the Golgi and plasma membrane is a consequence of further trafficking after initial insertion at the ER (Borgese et al., 2019). We selected mutant 1, a MAVS-R537A mutant of the mito TA construct (“mTA* protease”, Fig. S1) for our ER screen. The additional mutant constructs are described in Table S1.

**Figure S5.**
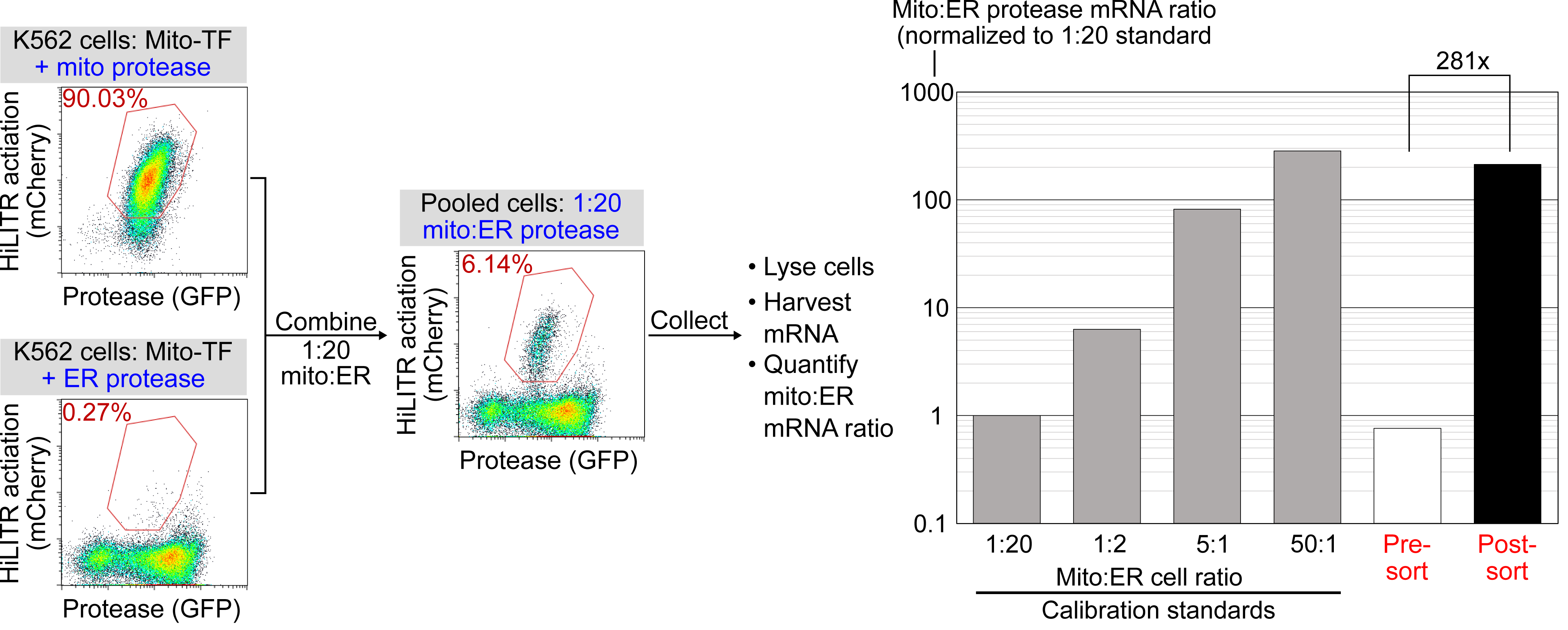
Model selection on K562 cells expressing mitochondrial TF HiLITR. Same as Figures 1E-F, except that cells expressing mitochondrial protease (co-localized with TF) are combined with cells expressing ER protease (rather than cytosolic protease as in Figure 1E). Cells were combined in a 1:20 ratio as indicated, stimulated with light for 3 minutes, and sorted for high mCherry expression 8 hours later. qPCR analysis of mito- and ER-protease transcript from predefined, pre-sort, and post-sort cell mixtures showed a 281-fold enrichment of mito-protease cells over ER-protease cells in one round of sorting.

**Figure S6.**
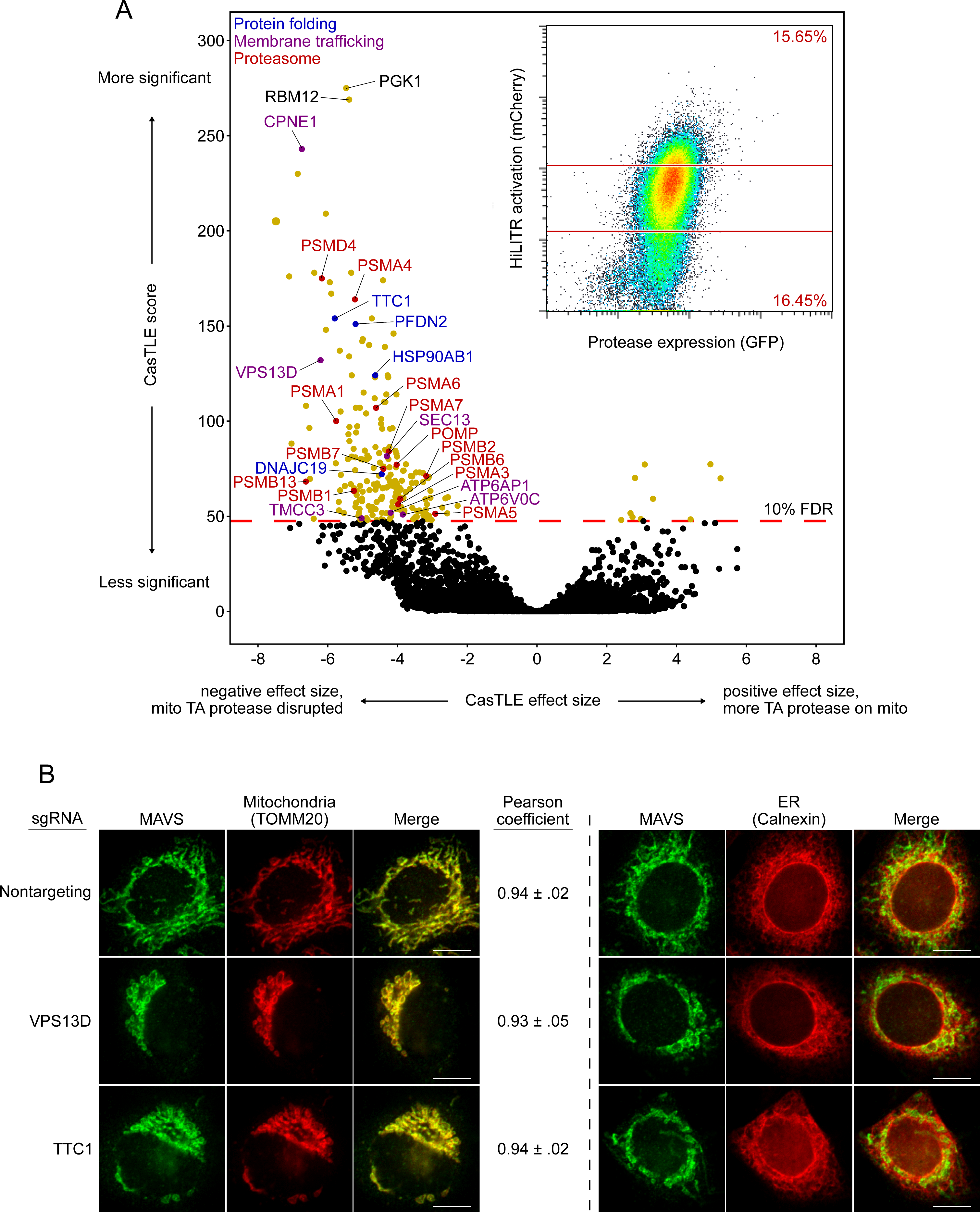
Whole-genome CRISPRi screen with HiLITR readout. A. CasTLE plot showing results of whole-genome CRISPRi screen, performed in two replicates in the clonal K562 mito TF/mito protease HiLITR cell line (shown in Figure 2C, left). FACS data in inset. X-axis shows log_2_-scaled change in HiLITR activation (ratio of high mCherry cells to low mCherry cells) relative to nontargeting controls. Y-axis shows the CasTLE score, a measure of significance. Labeled hits are annotated for function in protein folding (blue), membrane trafficking (purple), or are involved in proteasome function or biogenesis (red). B. Immunofluorescence microscopy of MAVS, an endogenous mitochondrial tail-anchored protein, with or without knockdown of two CRISPRi hits (VPS13D and TTC1) in HeLa. TOMM20 and Calnexin are mitochondrial and ER markers, respectively. Scale bar, 10 µm. For MAVS vs TOMM20 images, mean and standard deviation of the Pearson correlation coefficient between the protease and mito marker channels was calculated (n = ∼20 cells per condition).

## Text S6. Analysis of the whole-genome HiLITR screen. Related to Figure 2 and Figure S6

There are a number of interesting results to note from the whole-genome screen. First, the top hit, PGK1, is an artifact of using the PGK1 promoter upstream of HiLITR TF in the expression vector (the PGK1 promoter drives an antibiotic resistance gene, not the TF itself). Therefore, sgRNAs against PGK1 have off-target effects which silence the TF expression, producing a profound reduction in HiLITR activation. Interestingly, the second and third most significant hits, RBM12 and CPNE1, actually share a promoter (Yang et al., 2008). sgRNAs targeting RBM12 will knock down CPNE1, and vice versa, so it is encouraging that knockdown of RMB12 and CPNE1 produce very similar results.

Before moving to the three sublibrary screens (Fig. 2C-E), we performed follow-up on a few hits from the whole-genome screen. Two of these hits, VPS13D and TTC1, showed profound defects in mitochondrial morphology when the genes were knocked down by CRISPRi (Fig. S6B). VPS13D plays a role in organelle-to-organelle contact and bulk lipid transfer (Gao and Yang, 2018), and the effect of its depletion on mitochondrial morphology has previously been observed (Anding et al., 2018). TTC1 is a tetratricopeptide repeat (TPR) domain-containing protein that binds to both HSP70 (Liu et al., 1999) and HSP90 (Liou and Wang, 2005). Despite the general mitochondrial defects we observed, however, knockdown of VPS13D or TTC1 did not produce measurable changes in the colocalization of an endogenous mitochondrial TA protein (MAVS) with mitochondria (Fig. S6B).

After we performed the three sublibrary screens, we checked the results from VPS13D and TTC1 (Supplementary Table 2). While VPS13D has been observed to disrupt mitochondrial morphology (Anding et al., 2018), it has not been observed to disrupt the ER (Seong et al., 2018). Consistent with this, VPS13D knockdown decreased HiLITR activation in the TA and SA screens, but not the ER screen. Likewise, TTC1 knockdown decreased HiLITR activation in only the TA and SA screens. We speculate that VPS13D and TTC1 may have some functional relation.

**Figure S7.**
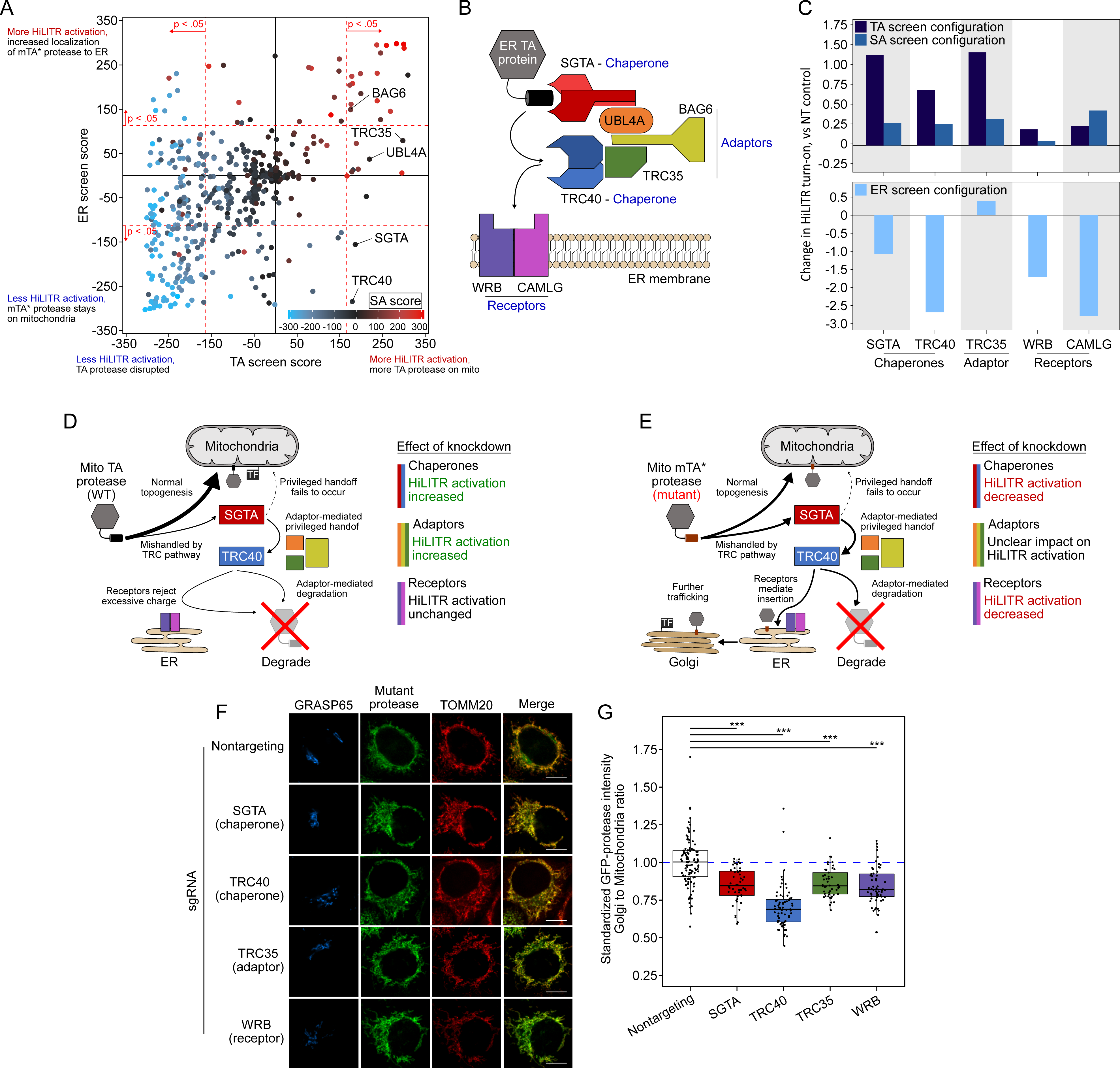
Re-testing of TRC/GET pathway genes with HiLITR. A. The plot from figure 3A, with genes in the TRC pathway (GET pathway in yeast) labeled. All 5 genes tested in the sublibrary screens (WRB and CAMLG were omitted) produced significant increases in HiLITR activity in the TA screen (p = 1.7e-7, hypergeometric test). B. Schematic of the TRC pathway. C. Quantitation of individual FACS analysis of gene knockdown in the TRC pathway. The K562 TA and SA (top) and ER (bottom) HiLiTR cell lines were transduced with individual sgRNAs against TRC pathway genes. Log_2_-transformed ratio of high mCherry to low mCherry cells was calculated for each plot and normalized to that of nontargeting (NT) control. D. Schematic showing possible membrane insertion pathway of TA protease. Most protein traffics to the OMM, but a subpopulation may be nonproductively handled by TRC pathway chaperones, resulting in rejection from ER insertion by the receptors, adaptor-mediated recruitment of ubiquitination machinery, and subsequent degradation. E. Schematic showing possible membrane insertion pathway of mTA* protease. F. Immunofluorescence microscopy analysis of TRC pathway knockdown. In HeLa cells, the localization of mTA* protease was compared to Golgi (GRASP65) and mitochondrial (TOMM20) markers. Scale bar, 10 µm. Note: knockdown of CAMLG in HeLa cells impaired cell adherence, preventing immunofluorescence analysis. G. Quantification of data in F, along with ∼20 additional fields of view per condition (total ∼50 cells per sample). For each cell, the mean intensity of Golgi-colocalized GFP was divided by the mean intensity of mitochondria-colocalized GFP. ***p < .001, Student’s T-test.

## Text S7. Performance of TRC pathway genes in the sublibrary screens. Related to figure S7

The TRC pathway is the first pathway discovered for the targeting of ER-destined tail-anchored proteins and is at this point well-characterized (Borgese et al., 2019). Figure S7B shows the key players in the TRC pathway. ER-targeted tail-anchored proteins that are TRC pathway clients are handled directly by two chaperones, SGTA and TRC40 (Chang et al., 2010; Schuldiner et al., 2008; Stefanovic and Hegde, 2007). Three adaptor proteins (UBL4A, TRC35, and BAG6) (Mariappan et al., 2010) coordinate handoff between SGTA and TRC40 (Shao et al., 2017; Wang et al., 2010), while BAG6 additionally recruits the E3 ubiquitin ligase RNF126 (not shown) to degrade nonproductively-associated proteins (Rodrigo-Brenni et al., 2014). At the ER membrane, WRB and CAMLG act as receptors to assist insertion of the client protein (Schuldiner et al., 2008; Vilardi et al., 2011; Yamamoto and Sakisaka, 2012), but proteins with significant positive charge flanking the transmembrane domain (such as mitochondrial tail-anchored proteins) are rejected, either by the receptors or due to the energetic barrier posed by the ER membrane (Rao et al., 2016).

The chaperones (SGTA and TRC40) and adaptors (TRC35, BAG6, and UBL4A) were included in the mito/ER sublibrary, while WRB and CAMLG were not. We further explored the TRC pathway with individual knockdown of the chaperones SGTA and TRC40, the adaptor TRC35, and the receptors WRB and CAMLG. Interestingly, the profiles of HiLITR performance across the three HiLITR configurations segregated based on the function of the protein in the TRC pathway (Fig. S7C). Notably, knockdown of none of the proteins affected the SA screen HiLITR configuration, consistent with the fact that the TRC pathway acts only on tail-anchored proteins.

In the TA screen configuration, knockdown of the chaperones and adaptors both increased HiLTR activation (Fig. S7A/C). If the chaperones are knocked down, there will be decreased mishandling of tail-anchored protein, and therefore an increase in normal topogenesis, localization to the mitochondria, and release of mitochondrial TF (Fig. S7D). Similarly, upon loss of adaptors, handoff of TA protein between SGTA and TRC40 is less coordinated. Adaptor mediated degradation of uninserted tail-anchored protein will also be reduced. Both such effects promote increased targeting of the tail-anchored protein to the mitochondrial membrane, increasing HiLITR activation (Fig. S7D). Knockdown of receptors in the TA screen configuration did not impact HiLITR activation (Fig. S7C). Since mitochondrial tail-anchored proteins that reach the ER are rejected on the basis of charge, loss of the receptors will have no additional impact (Fig. S7D).

In the ER screen configuration, knockdown of the chaperones and receptors decreased HiLITR activation (Fig. S7A/C). Loss of chaperones will mean less mTA* protease is routed to the ER, meaning less ends up colocalized with the ER-targeted TF (Fig. S7E). Similarly, if the receptors are disrupted, the mTA* protease cannot be inserted into the ER, decreasing HiLITR activation (Fig. S7E). In contrast to the chaperones and receptors, the adaptors gave mixed results, with two of the three adaptors producing no significant result in the ER screen (Fig. S7A/C). Knockdown of the adaptors will decrease handoff of the mTA* protease between SGTA and TRC40, providing opportunity for escape to the mitochondrial membrane (and decreased HiLITR activation). However, this effect is opposed by the fact that loss of adaptors will reduce degradation of the mTA* protease, providing more time for its insertion into the ER membrane (increasing HiLITR activity). As such, the impact of knocking down the adaptors is harder to predict for the ER screen (Fig. S7E).

Lastly, we tested the effect of TRC protein knockdown on the localization of the mutant tail-anchored protease. Knockdown of SGTA, TRC40, and WRB decreased activation of HiLITR in the ER screen configuration, indicating reduced ER-targeting of the mTA* protease. As expected, knockdown of any of these components reduced the fraction of mTA* protease colocalizing with the Golgi (Fig. S7F/G). TRC35 also decreased the fraction of mTA* protease colocalized with the Golgi, despite a neutral effect in the ER screen HiLITR configuration. It is likely that loss of TRC35 increases the fraction of mTA* protease which is rescued from the TRC pathway while also increasing the efficiency by which unrescued mTA* protease is inserted into the ER (S7E). The increase in mitochondrial mTA* protease and neutral effect on ER mTA* protease would decrease the *ratio* of Golgi-localized mTA* protease without affecting total Golgi-localized protease or subsequent HiLITR activation in the ER screen configuration.

**Figure S8.**
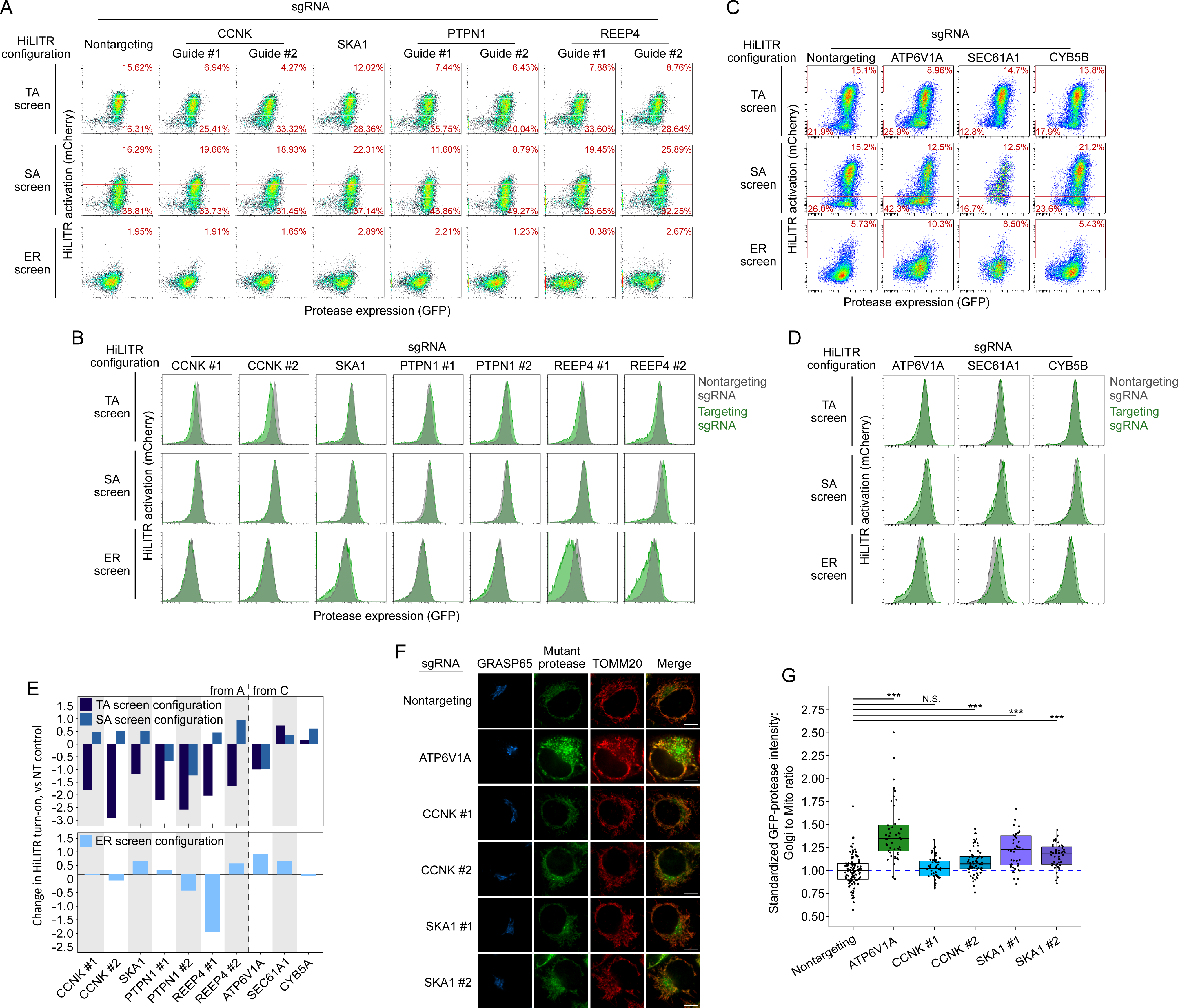
Re-testing individual sgRNAs from CRISPRi screen. A. FACS plots showing the effects of individual sgRNAs on HiLITR readout in three K562 HiLITR cell lines (TA, SA, and ER) from Figures 2C/D. Percentage of cells above and below the red lines shown in each plot. Genes in this group were hits with low TA score and mid to high SA score, labeled in Figure 3C. B. FACS plots showing the effects of the guides tested in (A) on protease expression level. Protease expression level in cells with nontargeting control sgRNA is overlaid in gray. C. Same as A, for three additional genes which were hits with low TA score and high ER score, labeled in Figure 3D. D. FACS plots showing the effects of the guides tested in (C) on protease expression level. Protease expression level in cells with nontargeting control sgRNA is overlaid in gray. E. Quantitation of FACS data in (A) and (C). Log_2_-transformed ratio of high mCherry to low mCherry cells was calculated for each plot and normalized to that of nontargeting (NT) control plot. Top shows TA configuration and SA configuration HiLITR data, bottom shows ER configuration HiLITR data for each gene tested. F. Fluorescence microscopy of mTA* protease in HeLa with knockdown of three different CRISPRi hits. GRASP65 and TOMM20 are Golgi and mitochondrial markers, respectively. Scale bars, 10 µm. H. Quantification of data in D, along with ∼20 additional fields of view per condition (∼50 cells per condition). For each cell, the mean intensity of Golgi-colocalized protease was divided by the mean intensity of mitochondria-colocalized protease. ***p < .001, Student’s T-test.

**Figure S9.**
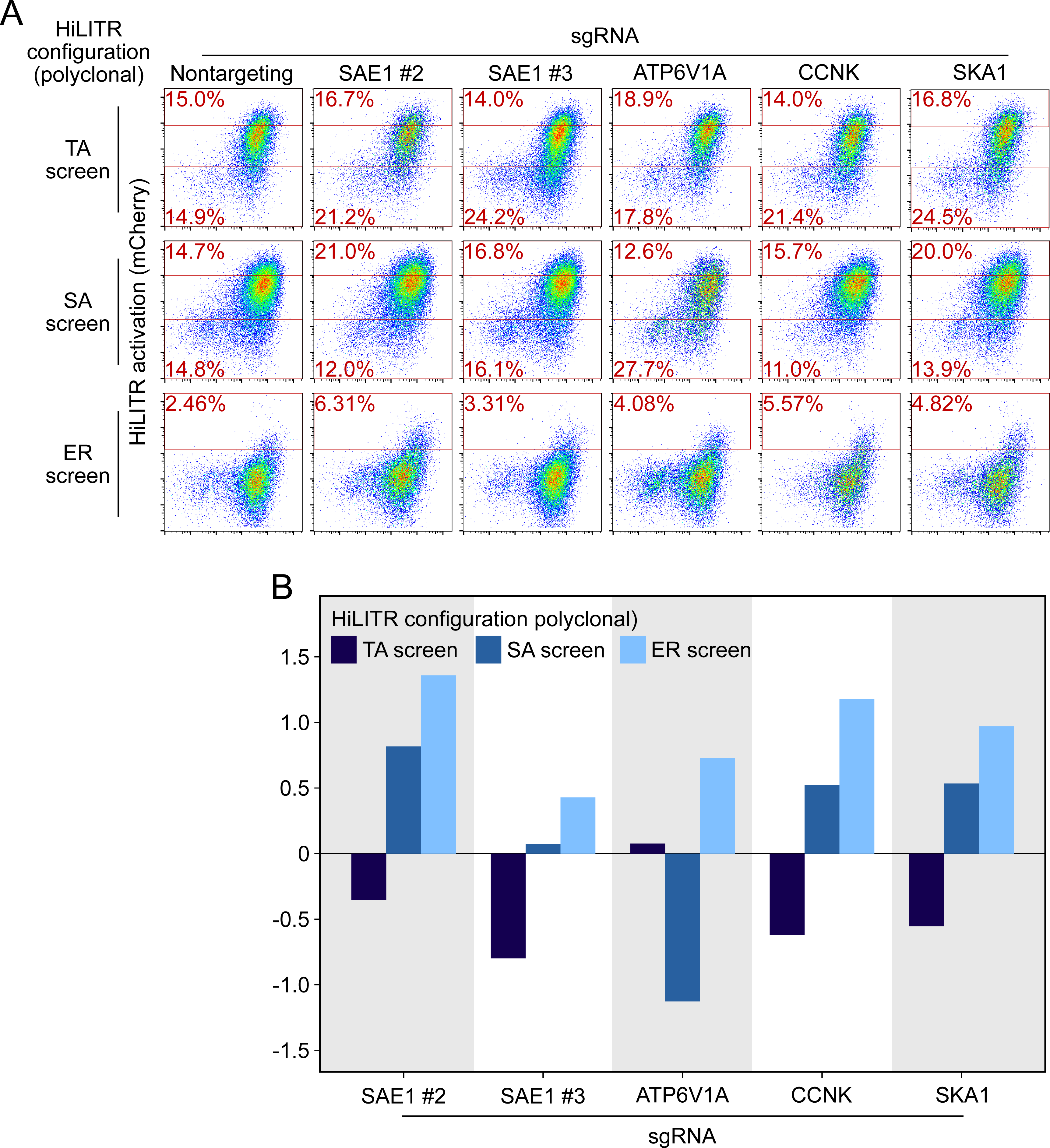
Re-testing individual sgRNAs in polyclonal cell lines. A. FACS plots showing the effects of individual sgRNAs on HiLITR readout in polyclonal K562 HiLITR cell lines (corresponding to TA, SA, and ER screens from Figure 2C/D). Percentage of cells above and below the red lines shown in each plot. B. Quantitation of FACS data in (A). Log_2_-transformed ratio of high mCherry to low mCherry cells was calculated for each plot and normalized to that of nontargeting (NT) control plot.

**Figure S10.**
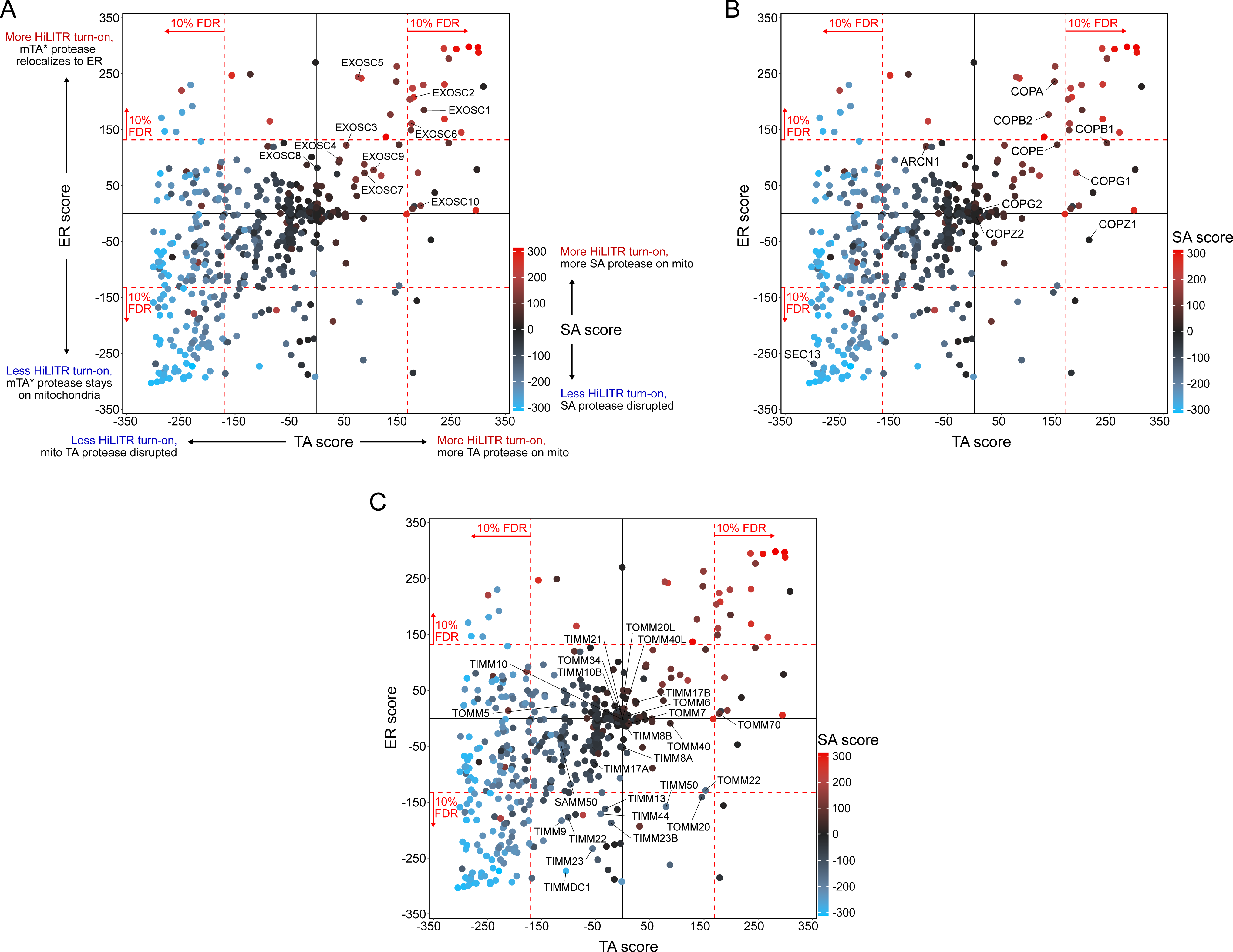
Other pathways of note. A. Results from the exosome complex. The plot from figure 3A, with genes in the exosome complex labeled. B. Results from the COPI/COPII pathway. The plot from figure 3A, with genes in the COPI pathway (Golgi to ER retrograde transport) and SEC13 from the COPII pathway (ER to Golgi anterograde transport) labeled. C. Results from TIMM and TOMM complexes. The plot from figure 3A, with the TIMM and TOMM complexes labeled.

## Text S10. Description of additional hits from Figures 3C-E and discussion of potential artifacts. Related to figures S8-S10

We performed individual validation experiments on a number of hits from sublibrary screens. Performance from individual validation experiments is compared to performance in the sublibrary screens in table S2.

PTPN1 is a tail-anchored protein known to localize to both the ER and mitochondria in an isoform-dependent manner (Brambillasca et al., 2006; Fueller et al., 2015). PTPN1 showed significant reduction in HiLITR activity in only the TA screen. We validated the strong negative effect of PTPN1 on HiLITR activation in the TA configuration with two guides (Fig. S8A/B/E). However, PTPN1 also decreased activation in the SA configuration, albeit to a more modest extent.

REEP4 plays a role in ER membrane sequestration during metaphase (Schlaitz et al., 2013). It was found to decrease HiLITR activation in the TA and ER screens, while increasing HiLITR activation in the SA screen. Validation of REEP4 with two guides was was consistent with screen results, with both guides showing decrease in activation in the TA configuration and one guide each showing either decreased activation in the ER configuration or increased activation in the SA configuration (Fig. S8A/B/E). REEP4 was observed to have a significant growth defect, and its annotation seemed to imply a nonspecific role, so we declined to analyze it further.

SEC61A1 is a member of the ER translocon. It was observed to decrease HiLITR activation in the TA and SA screens and increase HiLITR activation in the ER screen. During individual validation, it was instead observed to mildly increase HiLITR activation in all three configurations (Fig. S8C-E). Knockdown of SEC61A1 also produced a substantial growth defect.

CYB5B is a second tail-anchored protein known to localize to both the ER and mitochondria (D’Arrigo et al., 1993). It was seen to decrease HiLITR activation in the TA and SA screens and increase HiLITR activation in the ER screen (Fig. S8C-E). These results did not replicate with the guide we used for individual validation.

The proteins which replicated most successfully were SKA1, CCNK, and ATP6V1A (Fig. S12A-E). CCNK and SKA1 have annotation related to mitosis, while ATP6V1A is a member of the vacuolar ATPase. For these guides and for SAE1, we assessed the possibility that clone-specific effects could account for their performance in the HiLITR screens, as a consequence of either clone-specific sensitivity to certain biological pathways or off-target silencing if a HiLITR component was integrated next to a targeted gene. We performed HiLITR assays in the polyclonal cell lines from which the screen cell lines were isolated, and found that HiLITR performance in polyclonal cell lines was in general agreement with the results from the screens, though data were noisier due to the population heterogeneity (Fig. S9). ATP6V1A showed no loss of HiLITR activation in the heterogeneous TA configuration, while CCNK and SKA1 showed greater activation in the ER configuration than was observed with the clonal population.

In addition to individual hits, we took note of three groups of functionally related genes that showed patterns of activation in the sublibrary screens. Several components of the exosome complex increased HiLITR activation in one or more screens (Fig. S10A). This complex is almost certainly a false positive, as it plays a role in RNA degradation (Kilchert et al., 2016), but it importantly indicates that in some cases false positives can increase – rather than decrease – HiLITR activation. We noticed that the set of generally activity-increasing hits included several members of the COPI complex, which mediates ER-to-Golgi anterograde transport (Fig. S10B). Interestingly, SEC13, a member of the Golgi-to-ER retrograde COPII complex, decreased HiLITR activation in the TA and ER screen. While the mechanistic connection of COPI/II to trafficking of the HiLITR components is not readily apparent – and quite possibly indirect – the recapitulation of the opposing roles of COPI and COPII in the HiLITR data is intriguing. Finally, we looked at all proteins in the TOMM and TIMM complexes (Fig. S10C).

Surprisingly, knockdown of a large number of TIMM components, as well as TOMM20, decreased HiLITR activation in the ER screen. Several components also decreased HiLITR activation in the SA screen. We speculate that the knockdown of key TIMM and TOMM members might produce systems-level protein trafficking defects with far-ranging effects. It remains an important question how organelles in general, or different classes of proteins in specific, relate to global patterns in protein trafficking.

**Figure S11:**
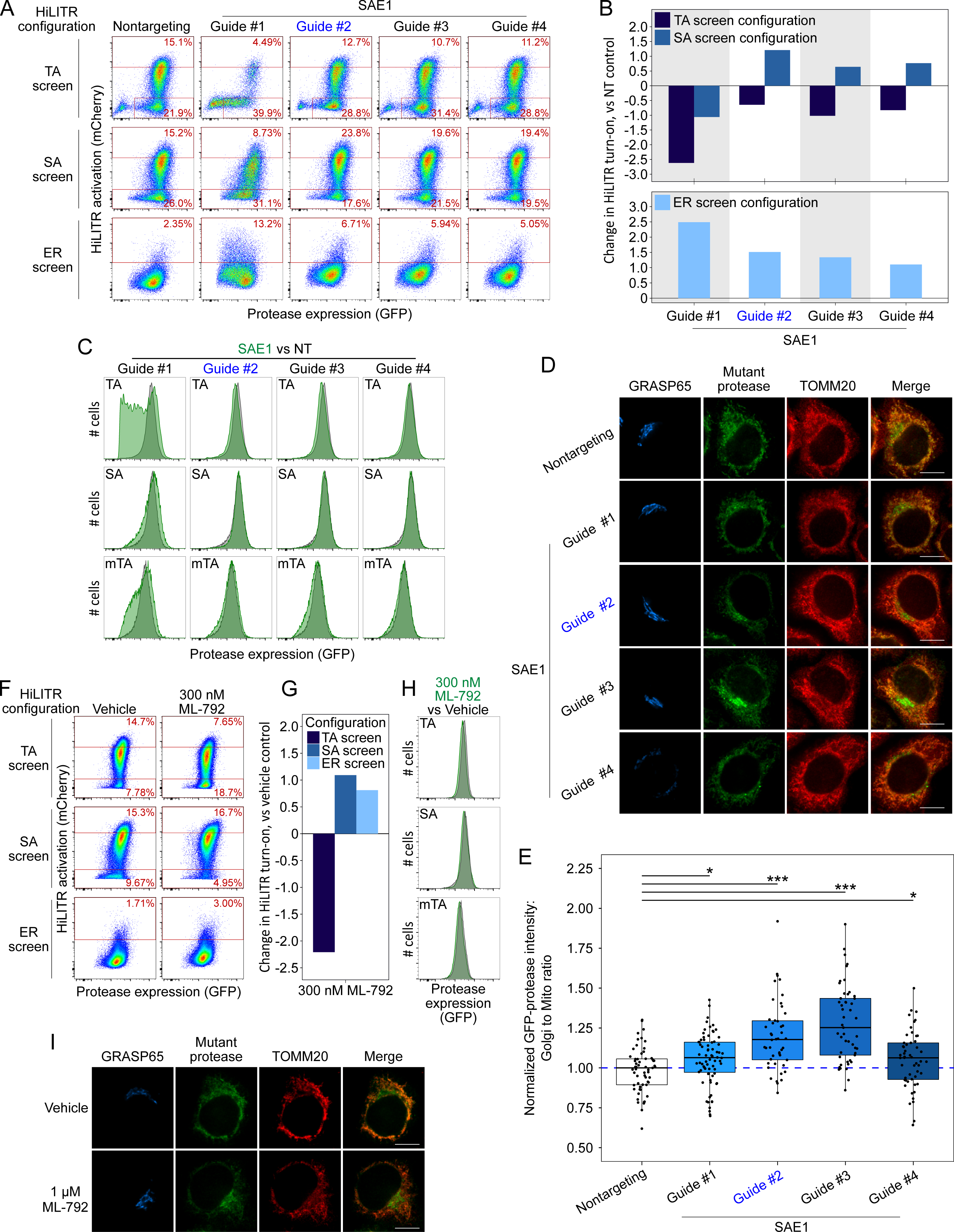
Additional data on SAE1. A. Same as Figure 4A, but showing data for three additional sgRNAs against SAE1. Guide #2 (blue) is the same guide used in Figure 4A, but with new data here. Note: Guide #1 produced a severe growth defect in the cells, potentially contributing to the discrepant HiLITR activation. B. Quantitation of FACS data in (A). Log_2_-transformed ratio of high mCherry to low mCherry cells was calculated for each plot and normalized to that of nontargeting (NT) control plot. Top shows TA configuration and SA configuration HiLITR data, bottom shows ER configuration HiLITR data for each gene tested. C. FACS analysis of GFP expression levels for the samples in (A). Green plot shows SAE1 knockdown cells; grey plot shows control cells (NT guide). D. Same as figure 4D, but with three additional sgRNAs against SAE1, and omitting ectopic construct expression. Guide #2 (blue) is the same guide as used in figure 4D. HeLa cells expressing mTA* protease and dCas9-KRAB were infected with SAE1 sgRNA or NT control for 8 days before expression of protease for 1 day, fixation, and imaging. Mitochondria and Golgi are visualized with anti-TOMM20 and anti-GRASP65 antibodies, respectively. Scale bar, 10 µm. E. Quantitation of data in (E) along with ∼20 additional fields of view (n = ∼50 cells per condition). For each cell, the mean intensity of Golgi-colocalized protease was divided by the mean intensity of mitochondria-colocalized protease. *p < 0.05, ***p < 0.001, Student’s t-test. F. FACS analysis of HiLITR activity upon chemical inhibition of SUMOylation. Three HiLITR configurations (as in Figures 2C/D) in K562 cells treated with either vehicle control (DMSO) or SUMO E1 ligase inhibitor ML-792 for two days prior to analysis. G. Quantitation of FACS data in (F). Log_2_-transformed ratio of high mCherry to low mCherry cells was calculated for each plot and normalized to that of vehicle control. H. FACS analysis of GFP-protease expression levels from the samples in (F) Green plot shows ML-792-treated cells, grey plot shows vehicle-treated control cells. I. Chemical inhibition of SUMOylation increases mislocalization of mutant tail-anchored protease to the Golgi. In HeLa cells, localization of the GFP-tagged mutant protease was compared to a Golgi marker (GRASP65) and a mitochondrial marker (TOMM20). Related to Figure 4I.

**Figure S12:**
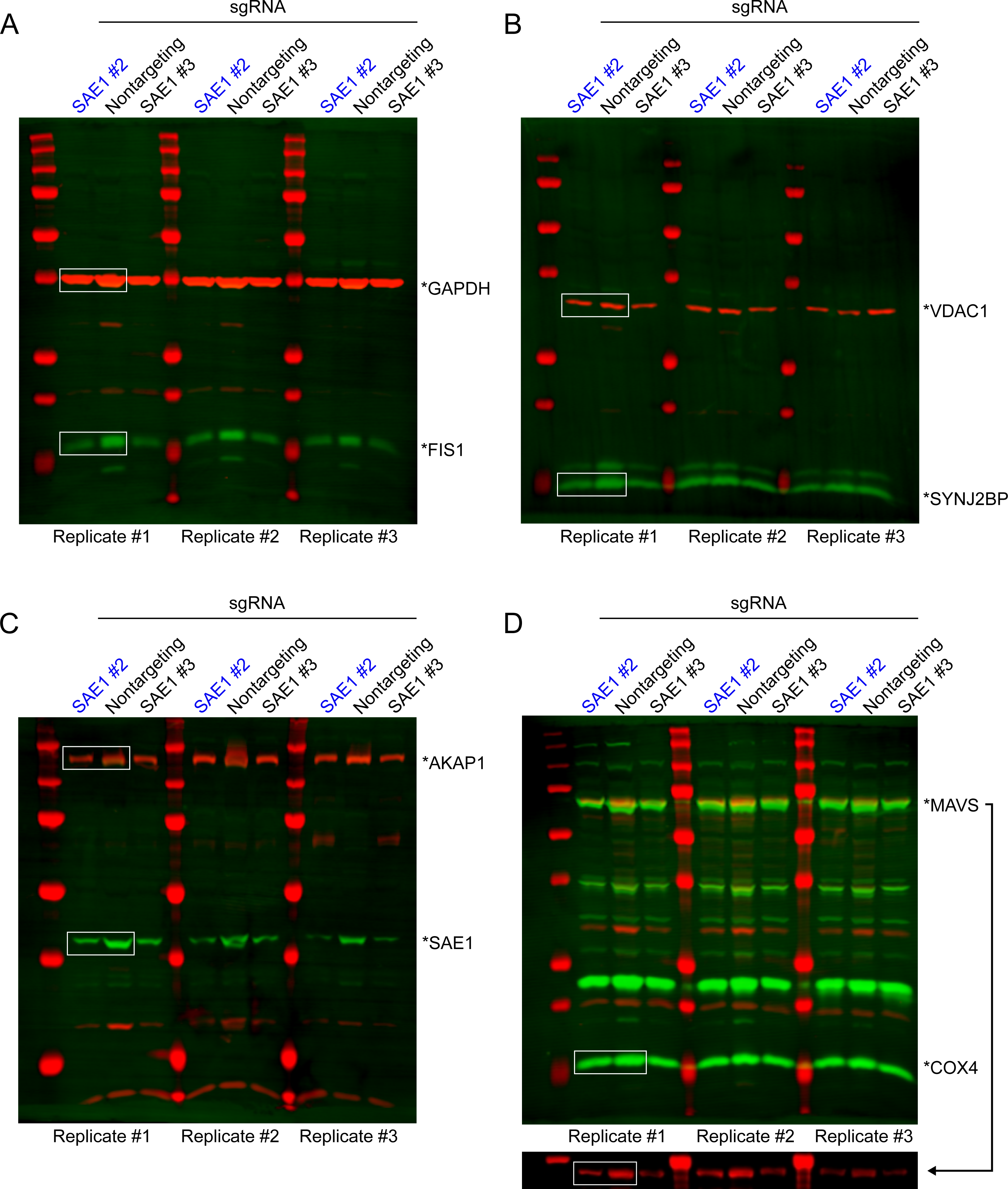
Uncropped western blots used to make Figure 4F/G. Data from sgRNA #2 (blue) was used to generate Figure 4F (boxed regions).

**Figure S13:**
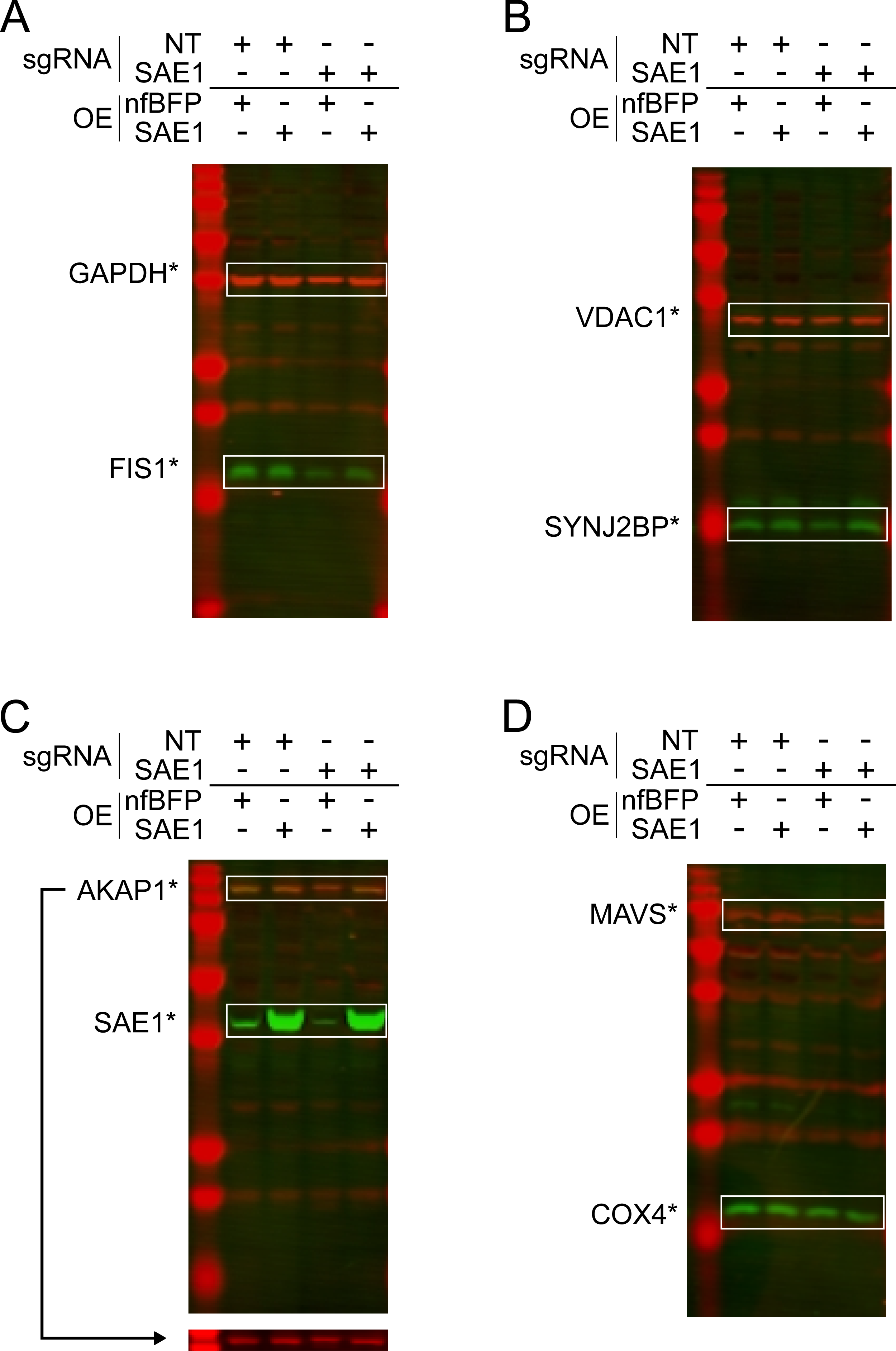
Uncropped western blots used to make Figure 4H. Data from sgRNA #2 (blue) was used to generate Figure 4H (boxed regions).

**Figure S14:**
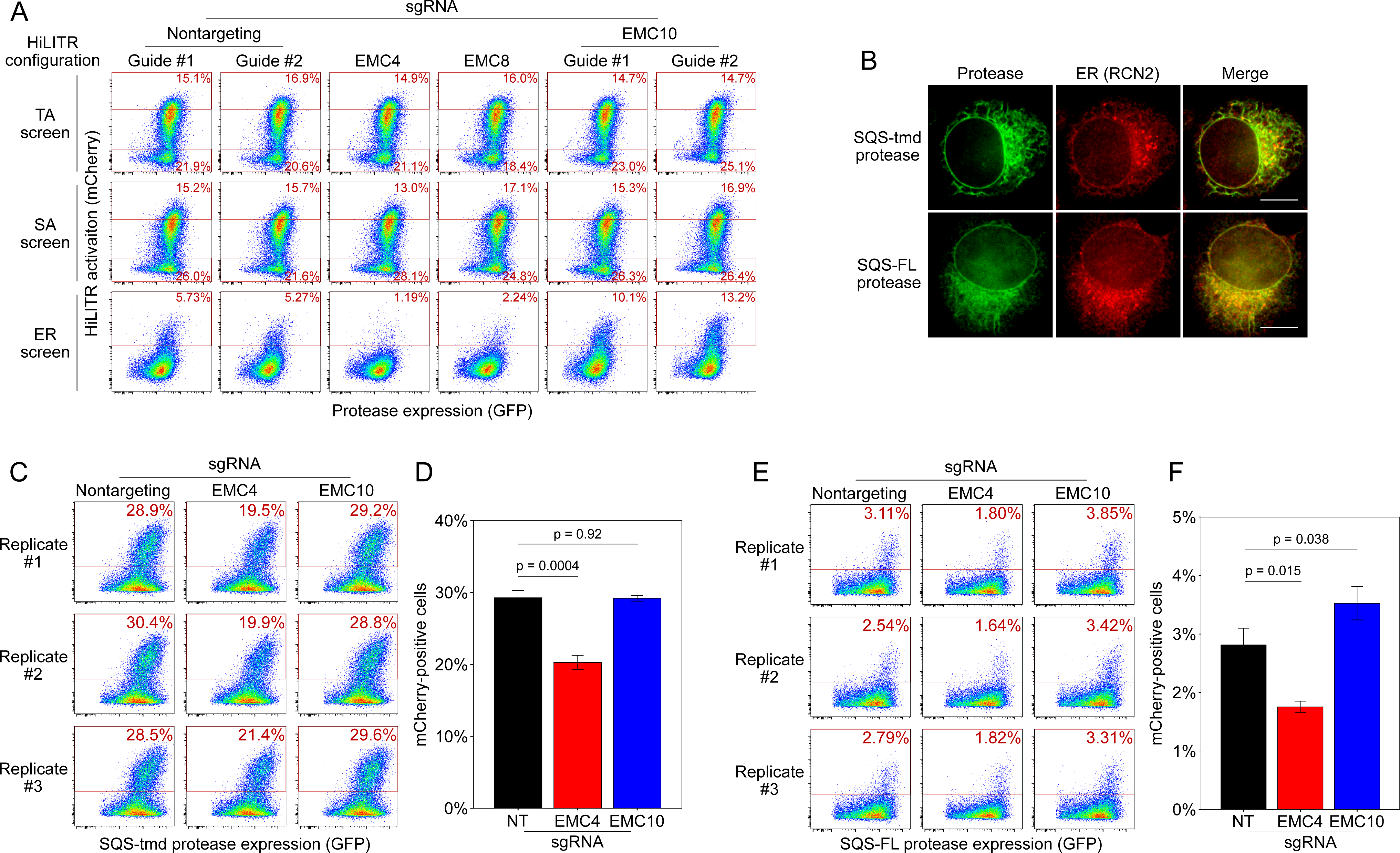
Additional HiLITR analysis related to the EMC. A. FACS analysis related to Figure 5B. Three HiLITR configurations (as in Figures 2C/D) in K562 cells with sgRNAs against EMC subunits and two nontargeting control guides. B. Immunofluorescence microscopy of the EMC client tail-anchored proteases. In HeLa cells, the localization of the SQS-tmd and SQS-FL proteases (GFP) were compared to an ER marker (RCN2). Scale bar, 10 µm. C. FACS analysis of HiLITR for ER-targeted tail-anchored protease. Protease was targeted to the ER with the transmembrane domain of TA protein SQS. HiLITR TF was also targeted to the ER. Three biological replicates were performed in polyclonal cell lines, and the percentage of cells above the red line is shown in each plot. D. Quantitation of the replicates in (B). Error bars = SEM. Significance calculated by Student’s t-test. E. Same as (B), but the protease was targeted to the ER with full-length SQS. F. Quantitation of the replicates in (D). Error bars = SEM. Significance calculated by Student’s t-test.

**Figure S15:**
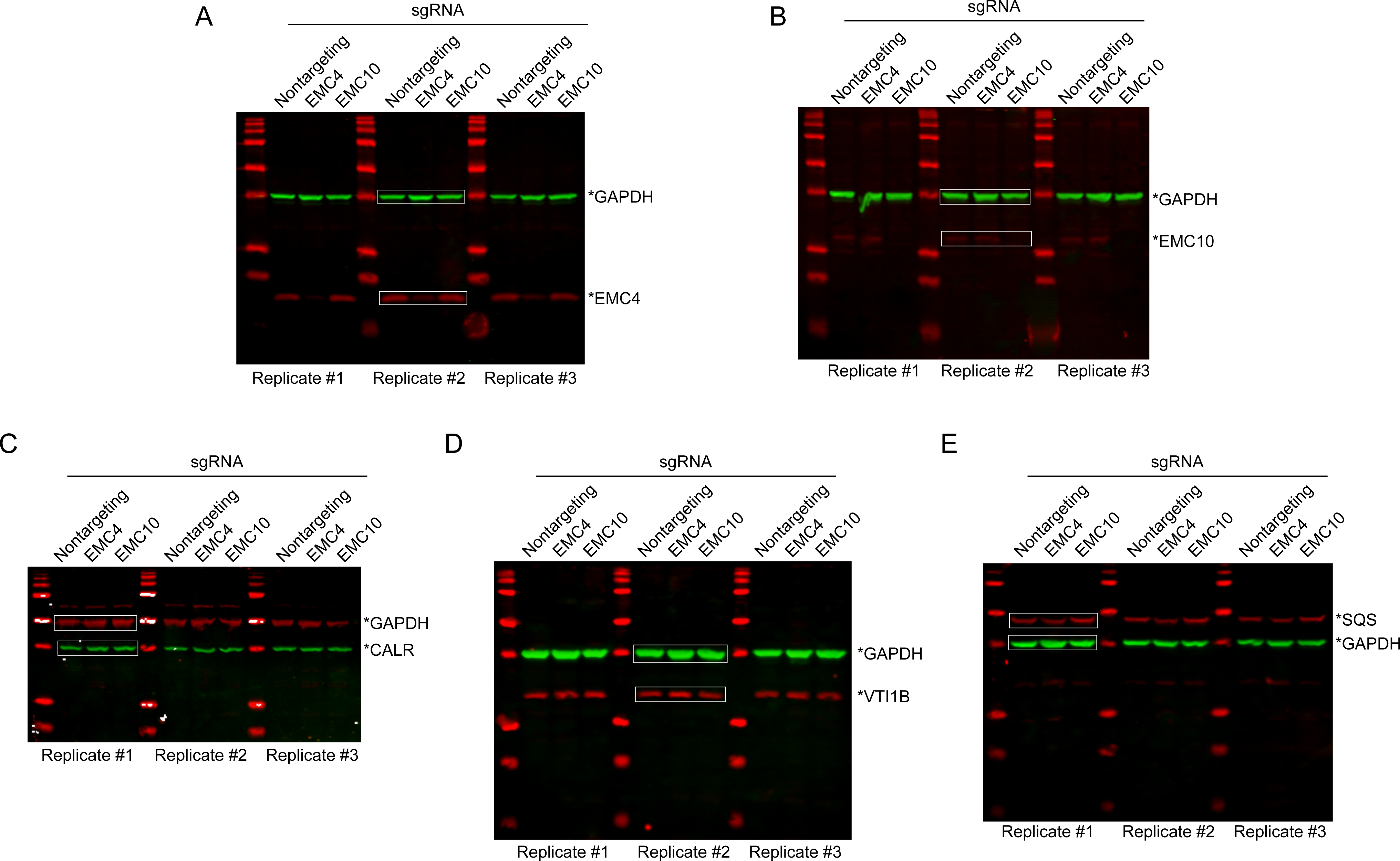
Uncropped western blots used to make. Figure 5E**/F.** Images in 5E were generated from the boxed regions in each plot, which are in each case the replicate most representative of the average shown in Figure 5F.

## Data S1

Plasmids used in the study and individual sgRNA sequences used.

- *Plasmids_Used*: Plasmid table for this study
- *sgRNAs_Used*: sgRNA sequences used for individual sgRNA sequences

## Data S2

Information about sgRNA libraries related to Figure S8 and Figure 3. Sequencing and CasTLE analysis results from the whole-genome screen and sublibrary screens. Comparison of individual validation data to sublibrary screen data.

- *WGS_sgRNAs*: sgRNA sequences and target genes in the whole-genome screen
- *TA_WGS*: CasTLE analysis of the whole-genome screen (TA screen HiLITR configuration)
- *Sublibrary_sgRNAs*: sgRNA sequences and target genes in the sublibrary screens
- *TA/SA/ER_Sublibrary*: CasTLE analysis of the sublibrary screens (TA/SA/ER screen HiLITR configurations)
- *Sublibrary_Comparison*: Comparison of combined-replicate CasTLE analysis across the TA/SA/ER sublibrary screens.
- *Hits&Validation*: Sublibrary screen data for genes mentioned in main and supplementary figures, with independent validation data appended where applicable.

